# An SNRNP70-eGFP knock-in zebrafish line reveals the physiological localisation and dynamics of endogenous SNRNP70 during development

**DOI:** 10.64898/2026.09.01.748523

**Authors:** Joshua Lloyd-Jones, Sophia Sartorius, David Gurevich, Nikolas Nikolaou

## Abstract

SNRNP70 is a core spliceosome RNA-binding protein best known for its essential role in nuclear pre-mRNA splicing. Although traditionally associated with nuclear RNA processing, previous studies have identified important extranuclear functions for SNRNP70 in neurons, including roles in mRNA stability, localisation, and axonal transport. Yet, much of our understanding of SNRNP70 localisation has relied on overexpression or transgenic approaches, leaving a critical gap in our knowledge of where endogenous SNRNP70 resides and how it behaves in living neurons under physiological expression conditions. Here, we address this limitation by establishing and validating a novel zebrafish SNRNP70-eGFP CRISPR knock-in line, enabling direct visualisation of the endogenous protein. eGFP was fused to the C-terminus of endogenous SNRNP70 while retaining the native 3′ untranslated region, preserving key regulatory features of the endogenous locus. We demonstrate that the knock-in faithfully reports endogenous SNRNP70 expression and reveals widespread physiological localisation throughout the developing nervous system, including prominent enrichment within axonal and synaptic compartments. Crucially, live *in vivo* imaging reveals that endogenous SNRNP70 is dynamically localised within neuronal mRNP granules, providing direct evidence of its physiological behaviour in these structures without the confounding effects of protein overexpression. Proximity ligation analyses further demonstrates associations between endogenous SNRNP70 and PABPC1B, FUS, and UPF1, which are established neuronal mRNP granule components. Together, our work provides a validated genetic and imaging resource for investigating SNRNP70 at endogenous levels in the living nervous system. By overcoming key limitations of conventional transgenic and overexpression-based approaches, the SNRNP70-eGFP knock-in enables physiological analysis of SNRNP70 localisation and dynamics and reveals its prominent and dynamic organisation within neuronal mRNP granules. More broadly, this work highlights how endogenous fluorescent tagging can provide a versatile platform for resolving the spatial organisation of RNA-binding proteins in living neurons under physiological expression conditions and provide new insight into the regulation of neuronal mRNA fate.

## Introduction

SNRNP70 is a core component of the spliceosome and an essential RNA-binding protein (RBP) involved in pre-mRNA splicing (Kondo et al., 2015). As a component of the U1 small nuclear ribonucleoprotein (snRNP), SNRNP70 recognises and binds the U1 small nuclear RNA and contributes to spliceosome assembly and intron recognition. Its canonical function is therefore closely linked to the nucleus and to the early stages of gene expression, where pre-mRNA processing determines which transcripts are competent for subsequent export and translation.

Neurons are highly polarised cells in which the nucleus-containing soma can be separated from distal axons, dendrites and synaptic terminals by considerable distances. Consequently, neuronal gene regulation extends well beyond the nucleus, with RNA transport, local mRNA stability and spatially restricted translation providing mechanisms through which gene expression can be regulated within individual neuronal compartments (Holt et al., 2019). These processes are coordinated by dynamic messenger ribonucleoprotein (mRNP) complexes containing mRNAs and RBPs, which can regulate transcript transport, storage, translation and decay. In neuronal processes, such complexes can assemble into dynamic mRNP granules that respond to developmental and extracellular signals, including cues that regulate axonal growth and synaptic function (Banani et al., 2017; Kedersha et al., 2005; Koppers et al., 2019; Liao et al., 2019; Welshhans & Bassell, 2011).

Although traditionally considered a predominantly nuclear splicing factor, SNRNP70 has also been detected in extranuclear neuronal compartments (Nikolaou et al., 2022; Thomas-Jinu et al., 2017). We previously demonstrated that cytoplasmic SNRNP70 is required for motor neuron connectivity in zebrafish and found that cytoplasmic SNRNP70 influences the stability, localisation and axonal transport of mRNAs (Nikolaou et al., 2022). SNRNP70 has also been reported to localise to neuronal cytoplasmic compartments associated with mRNA targets, providing further evidence that its function may extend beyond the nucleus (Nikolaou et al., 2022; Thomas-Jinu et al., 2017). Together, these observations highlight the need for a tool that enables direct visualisation of endogenous SNRNP70 in a living nervous system and under physiological expression conditions.

Addressing this question requires visualisation of endogenous SNRNP70 within its native cellular context. Conventional approaches, including overexpression of fluorescently tagged proteins and transgenesis, can provide valuable insights into subcellular localisation but may not faithfully recapitulate endogenous protein abundance, spatial expression, or temporal regulation. Elevated protein levels may themselves alter subcellular localisation or promote molecular interactions that are not representative of endogenous SNRNP70 behaviour (Maharana et al., 2018). Furthermore, because transgenes are removed from their native genomic environment, they may lack regulatory elements that contribute to physiological expression patterns. This is particularly relevant for SNRNP70, whose expression and subcellular distribution may be regulated in a developmentally and cell-type-specific manner (Nakaya et al., 2013; Rösel-Hillgärtner et al., 2013). An endogenous fluorescent reporter therefore provides a means of distinguishing physiological SNRNP70 localisation and distribution from patterns arising from altered protein abundance or ectopic expression, while enabling the same protein to be tracked across developmental stages and neuronal compartments.

CRISPR-mediated knock-in provides a means of overcoming these limitations by fluorescently tagging the endogenous protein at its native genomic locus (Levic et al., 2021; Zhang et al., 2023). Such an approach preserves endogenous transcriptional regulation and, depending on the targeting strategy, can retain regulatory sequences within the native transcript. An endogenous fluorescently tagged SNRNP70 therefore provides an opportunity not only to determine where SNRNP70 is expressed, but also to examine its distribution, dynamics and subcellular organisation in living developing neurons under physiologically relevant expression conditions.

Here, we established and validated a novel zebrafish SNRNP70-eGFP knock-in line in which eGFP is fused to the C-terminus of the endogenous SNRNP70 protein while preserving the endogenous 3′ untranslated region (3′UTR). We demonstrate that the knock-in line maintains *snrnp70* transcript expression, produces the expected SNRNP70-eGFP fusion protein, and preserves the major features of endogenous SNRNP70 expression and localisation, including its predominant nuclear and extranuclear pools. Using this endogenous reporter, we reveal a widespread distribution of SNRNP70 throughout the developing nervous system, including prominent localisation within neuronal processes and punctate cytoplasmic structures. Live imaging further demonstrates that endogenous SNRNP70 is associated with dynamic, motile mRNP granules *in vivo*, extending previous observations based on transgenic SNRNP70 expression. Finally, proximity ligation analysis demonstrates the utility of the endogenous reporter as a physiologically relevant model for investigating molecular associations between SNRNP70 and neuronal RNA-regulatory proteins. Together, these findings establish the SNRNP70-eGFP knock-in as a robust resource for visualising and investigating endogenous SNRNP70, while providing new evidence that SNRNP70 is spatially organised within dynamic neuronal mRNP granules and associated with the RNA regulatory machinery beyond its canonical nuclear splicing role.

## Materials and methods

### Animals

Adult zebrafish were reared at 28.5°C on a 14 h light/10 h dark cycle. Embryos were obtained from in-house breeding zebrafish stocks by pairwise light-induced spawning, collected for microinjection and raised in E3 media (5 mM NaCl, 0.17 mM KCI, 0.33 mM CaCl_2_, 0.33 mM MgSO_4_) in an incubator at 28.5°C until needed in experiments. Two new genetically altered lines were generated and used here: TgKI(*snrnp70-eGFP*)*^ba9^* and Tg(*mnx1:mCherry*)*^ba126^*. For the TgKI(*snrnp70-eGFP*)*^ba9^*line used in this paper, the F_1_ generation was created by outcrossing with the *mitfa^w2^* fish (Lister et al., 1999). This work was approved by the local Animal Welfare and Ethical Review Bodies (University of Bath and University of Exeter) and was caried out in accordance with the Animals (Scientific Procedures) Act, 1986, under project license (PP8698401) from the United Kingdom Home Office.

### Generation of TgKI(*snrnp70-eGFP*) line

The TgKI(*snrnp70-eGFP*) line was established by following a previously published CRISPR/Cas9 approach (Levic et al., 2021). eGFP was inserted at the C-terminal downstream of exon 10. Homology arms were used to ensure specificity (see below); one homology arm being composed of the entire 3’UTR region of the endogenous zebrafish *snrnp70* gene.

#### Generation of the donor plasmid

A fragment of ≈1.3kb, spanning from 320bp upstream to the 3’ end of intron 8 to just prior to the stop codon in Exon 10, was amplified from the s*nrnp70* locus and subcloned into the *pUC19-TgKI-MCS-eGFP-MCS-polyA* vector using the primers s*nrnp70*-EcoRI-forward 5’-GATGAATTCTGGCTTCTGTAGTTTGGTTA-3’ and *snrnp70*-XbaI-reverse 5’-ATTAGAGGTGGCGGTGGCGACCGGCCGGTGGATCCTGTACTCATCACCCTGGGCCT-3’, using the restriction enzymes EcoRI and XbaI. The entire 3’UTR of the zebrafish s*nrnp70* gene was amplified and subcloned into the resultant vector using the primers 3’UTR-HindIII-forward 5’-CCGAAGCTTAAGTGCCATCGTTGTGTGAATGATT-3’ and 3’ UTR-AflII-reverse 5’-CGGACATGTCCGTTCGGCAATAAAAAAGTAT-3’. The gRNA target site was mutated in this vector using site-directed mutagenesis using the primers gRNA-SDM-forward 5’-GGGATAGGCAAGGTCTTCGATGCCCACAGTT-3’ and gRNA-SDM-reverse 5’-AACTGTGGGCATCGAAGACCTTGCCTATCCC-3’

#### Preparation of the injection mixture, microinjection and screening of zebrafish embryos

10 µM crRNA, 20 µM tracrRNA, 2 µM recombinant Cas9 protein and 10 ng/µL donor plasmid were diluted in nuclease free water with 10% Phenol red and left at room temperature until egg collection and injections. 1.5-1.8 nl of injection mix was injected directly into the cell or the yolk just beneath the cell at the one-cell stage zebrafish embryos. Injected embryos were reared to 24hpf, following which they were screened for eGFP fluorescence under a Leica MZFLII Fluorescence Stereomicroscope. Only those embryos that displayed eGFP fluorescence were selected for genomic DNA extraction and sequencing, or for rearing as potential founders.

#### gDNA sequencing of F_0_ founder offspring

To confirm the insertion of eGFP into the C-terminus of SNRNP70, gDNA was extracted from F_1_ offspring positive for eGFP fluorescence. Zebrafish were anaesthetised in a Tricaine solution, and the dorsal lobe of the caudal fin was removed using a sterile, sharp, scalpel blade. The sample was transferred into a sterile 1.5 mL tube and stored on ice. The gDNA was then extracted by resuspending the fin in 50 µL of Lysis Buffer (10 mM Tris HCl pH 8.0, 50 mM KCl, 0.3% Tween-20, 0.3% IGEPAL, 1 mM EDTA, 2 mg/mL Proteinase K (ThermoFisher Scientific, AM2546), following which the samples were incubated overnight at 56°C. The Proteinase K was then deactivated by heating at 98°C for 10 minutes and the samples were stored at -20°C until further use. The regions surrounding the integration site were sequenced using the primers Genotyping-forward 5’-TGGCTTCTGTAGTTTGGTTAGGAATG-3’ and eGFP-N-terminal-reverse 5’-AGCTCCTCGCCCTTGCTCAC-3’ to cover the sequence upstream of the gRNA site and into eGFP; eGFP-C-terminal-forward 5’-GGCATGGACGAGCTGTACAAGTAA-3’ and *lin7b*-reverse 5’-ACCACACGTCACAGGCGCGA-3’ to sequence from eGFP to the neighbouring gene *lin7b*.

### Generation of Tg(*mnx1:mCherry*) line

To establish a Tg(*mnx1:mCherry*) founder generation, *pBSIISK-Scel-mnx1-mCherry-polyA* plasmid (See et al., 2014) was injected into fertilised eggs at one-cell stage together with I-SceI magenuclease according to previous methods (Grabher et al., 2004). Founder fish were then screened for germline transmission into F_1_ generation, and a stable zebrafish line was established.

### Immunohistochemistry-Immunofluorescence (IHC-IF)

#### Sample preparation

Zebrafish larvae were fixed in a 4% paraformaldehyde solution for 2 hours and 30 minutes at room temperature. Fixed samples were then embedded in a noble agar-sucrose solution (0.5% agar, 1.5% sucrose) in PBS, and set overnight at 4°C. The sample-embedded agar blocks were removed from the mould and placed in a 30% sucrose/PBS solution overnight at 4°C. Sample blocks were then removed from the sucrose solution and embedded in OCT matrix on a cryosectioning chuck and snap frozen in dry ice. Transverse sections of 30 μm thickness were cut along the rostral-caudal axis of the larvae, using the Leica CM1850 cryostat. Sample sections were mounted on SuperFrost slides, with samples set in a boundary by a hydrophobic marker.

#### Immunohistochemistry-Immunofluorescence

For immunohistochemistry of cryosectioned tissue, the samples were taken through the following protocol. Sections were washed once with PBS, for 10 minutes, following which they were permeabilised by incubating in a 0.1% PBST (Triton X-100) solution for 30 minutes at room temperature. Sections were then blocked with a 10% goat serum/PBST solution for 1 hour at room temperature. The sections were then incubated overnight at 4°C in a 10% goat serum/PBST solution containing the primary antibodies. The following day, sections were washed three times in PBST, for 15 minutes, following which they were incubated for 1 hour at room temperature in a 10% goat serum/PBST solution containing the Alexa-Fluor conjugated secondary antibodies. Sections were then washed three times, for 15 minutes each, following which they were incubated in a 1.25 μg/mL DAPI in PBST solution for 10 minutes, and then washed briefly before mounting in FluorSave Mounting Reagent and covered by a glass slide. The primary antibodies and their dilutions: chick anti-GFP 1:500 (Fisher Scientific, PA19533); rabbit anti-PABPC1B 1:250 (GeneTex, GTX128930); anti-FUS 1:200 (Affinity Bioscience, DF8391); anti-UPF1 1:300 (Stratech/ DF6440-AFF); and mouse anti-acetylated α-Tubulin 1:500 (Sigma-Aldrich, T7451). The secondary antibodies and their dilutions (all used at 1:500): Goat anti-chick IgG (H + L) Alexa FluorTM-488 (ThermoFisher Scientific, A11039), Goat anti-rabbit IgG (H + L) Alexa FluorTM-568 (ThermoFisher Scientific, A21069), Goat anti-mouse IgG (H + L) Alexa FluorTM-633 (ThermoFisher Scientific, A21050).

### Zebrafish primary neuronal cell culture

Zebrafish embryonic primary neuron cultures were carried out as previously published (Taylor and Houart, 2024; Sassen et al., 2017) with minor modifications. Briefly, following pair-mating and collection, embryos of the TgKI(*snrnp70-eGFP*) line were incubated in Autoclaved Fish Water (AFW; E3 medium, 1X Penicillin-Streptomycin (Fisher Scientific, 11568876), 50 μg/mL Gentamycin (Fisher Scientific, 12664735), 0.001% methylene blue (Fisher Scientific, 4841-16)) until 24 hpf, at which point they were screened for green fluorescence on a Leica MZFLII Fluorescence Microscope and 400 eGFP^+^ embryos selected. Embryos were then transferred into a sterile 40 μM cell strainer (Fisher Scientific, 11587522), and the outer chorions sterilised by incubation in a 0.003% sodium hypochlorite (Fisher Scientific, S/5042/15) in AFW solution for 5 minutes, followed by 5 minutes in AFW, 5 minutes in the 0.003% sodium hypochlorite/AFW, and a final 5 minutes in AFW. Embryos were then enzymatically dechorionated via incubation in a 0.5 mg/mL Pronase (Merck, 1016592100)/AFW solution for 15 minutes, followed by several washes in AFW to remove the chorions. Embryos were then transferred into a clean Petri dish, placed inside a sterile cell strainer, and moved to tissue culture. In the cell culture hood, embryos were sterilised by incubation in a 35 mm Petri dish containing a 70% Ethanol solution for 5 seconds, followed by a wash in a separate petri containing L-15 medium (Fisher Scientific, 15460554). The embryos were then transferred into a sterile 0.5ml tube, washed twice with HBSS (Fisher Scientific, 11580466), and then enzymatically dissociated in 1mL of a 0.25% Trypsin (Fisher Scientific, 11538876)/HBSS solution for 15-25 minutes with gentle trituration with a P1000 pipette throughout. Once large fragments of tissue were no longer visible, the cell suspension was filtered through a 40 μM strainer into a conical 50mL Falcon Tube (Merck, 227261), and the volume topped up with 2 ml of Zebrafish Neural Medium (L-15, 1X MACS Neurobrew (Miltenyi Biotec, 130-093-566), 1X N-2 Supplement (Fisher Scientific, 17502048), 2% Heat Inactivated Foetal Bovine Serum (FBS; Merck, F9665-50ML), 10 ng/mL BDNF (PeproTech, 450-02), 1X PenStrep, 50 μg/mL Gentamycin). The cells were then pelleted by centrifugation at 300 g for 7 minutes at 4°C, following which the excess media was removed, and the cell pellet resuspended in ZNM, at a ratio of 1 ml ZNM:400 embryos. The cell suspension was then plated, at a volume of 60 μL per well, into a 24-well plate (Fisher Scientific, 10624903) containing 12 mm glass coverslips (VWR, 631-1577) which had been coated with 0.1 mg/mL Poly-D-Lysine (Fisher Scientific-16021412) and 4 μg/mL laminin (Merck, 11243217001) solutions. The plate was then wrapped in parafilm and incubated at room temperature until 12 days in vitro (DIV), after which the coverslips were used for experimental assays.

### Immunocytochemistry-Immunofluorescence (ICC-IF)

At 12DIV, TgKI(*snrnp70-eGFP*) neuronal cell cultures were fixed by removing the ZNM, washing once with Dulbecco’s Phosphate Buffered Saline (dPBS; Fisher Scientific, 14190094), and incubating in a 4% paraformaldehyde (PFA) solution (Fisher Scientific, 15434459) for 10 minutes. The cells were then washed once with dPBS and then incubated in 1mL ice-cold 100% methanol for 10 minutes at room temperature. The methanol solution was then removed, samples washed three times with dPBS, and the cells were then permeabilised for 15 minutes at room temperature in 1mL of a 0.1% Triton X-100 PBS (PBST) solution. Following this, coverslips were then blocked in a 5% Donkey Serum (DS; Abcam, ab7475)/0.1% PBST solution for 30 minutes at room temperature, after a 1% DS/0.1% PBST solution containing the primary antibodies was added, and the cells incubated overnight at 4°C. The primary antibody solution was removed, the coverslips were then washed three times, for five minutes each, with 0.1% PBST. Following this the coverslips were incubated in a 1% DS/0.1% PBST solution containing the secondary antibodies, overnight at 4°C.The secondary antibody solution was removed, and the coverslips washed three times, for five minutes each, with 0.1% PBST. After the final wash, the coverslips were incubated in a 1.25 μg/mL DAPI/H_2_O solution for 1 minute, followed by two final dPBS washes. The coverslips were then mounted on a SuperFrost Plus slide (VWR, 631-9483) on a small droplet of FluorSave Mounting Reagent (Merck, 345789-20ML). The primary antibodies and their dilutions, along with the secondary antibodies and their dilutions, were the same as with the IHC-IF of SNRNP70eGFP KI embryos.

### Proximity ligation assay (PLA)

At 12DIV, cell culture coverslips were fixed using the same method as the ICC-IF. Following fixation, the coverslips were incubated in Quenching solution (50 mM Ammonium Chloride (NH_4_Cl); Fisher Scientific-1139173) for 15 minutes at room temperature. This solution was removed, the coverslips washed once with dPBS, following which they were permeabilised in 0.1% PBST for 15 minutes at room temperature. After permeabilisation, coverslips were washed three times with dPBS. At this point, the coverslips were taken through the proximity ligation assay, following the manufacturer’s instructions. For the purposes of this assay, the blocking step was conducted in the 24-well plate. Following this, the coverslips were moved from the 24-well plate into a humidity chamber (made by covering a 140 mm Petri Dish (Cromwell, STS3855006F) in foil and lining the inner rim with damp tissue paper) and incubated in the primary antibody solution in a 37°C incubator for 1 hour. For the washes following the primary antibody incubation, the coverslips were moved back into a 24-well plate. For the ligation reaction, the coverslips were then moved into the humidity chamber and incubated at 37°C for 30 minutes. The coverslips were then washed again after transferring to the 24-well, following which the amplification step was conducted in the humidity chamber for 1 hour and 40 minutes at 37°C. After the amplification step, the coverslips were transferred from the humidity chamber to a 24-well for the final washes. Following the final wash, the coverslips were incubated in a primary antibody containing solution (1% DS/0.2% PBST) overnight at 4°C to stain for acetylated α-Tubulin. The solution was removed, and the coverslips washed three times with dPBS, for five minutes each, after which the coverslips were incubated in a secondary antibody containing solution (1% DS/0.2% PBST) for 1 hour at room temperature. Following this, the solution was removed and the coverslips washed three times with dPBS, for five minutes each, after which the coverslips were incubated in a 1.25 μg/mL DAPI solution for one minute. The coverslips were then washed twice, for five minutes, with dPBS, after which they were mounted in a drop of FluorSave media on a SuperFrost Plus slide. The primary antibodies used for the proximity ligation assay, and their dilutions: goat anti-GFP 1:500 (Abcam, ab6673); rabbit anti-PABPC1B 1:250 (GeneTex, GTX128930); anti-FUS 1:200 (Affinity Bioscience, DF8391); rabbit anti-UPF1 1:300 (Stratech, F6440-AFF). The Duolink proximity ligation probes: Duolink Anti Goat MINUS (Merck, DUO92006-30RXN); Duolink Anti Rabbit PLUS (Merck, DUO92002-100RXN). The primary antibody used for the ICC-IF: Mouse anti-acetylated α-Tubulin 1:500 (Merck, T7451-200UL). The secondary antibody used for the ICC-IF: Goat anti-mouse IgG (H + L) 1:500 Alexa FluorTM-488 (ThermoFisher Scientific, A11001).

### Subcellular fractionation

At 96 hpf, 150-200 eGFP^+^ embryos from TgKI(*snrnp70-eGFP*) line were anaesthetised and enzymatically dechorionated via incubation in a 0.5 mg/mL Pronase solution, for 15 minutes. The dechorionated embryos were washed briefly with E3 solution, transferred into a 0.5 mL tube, and the excess solution removed. The weight of the embryos within the tube was measured on an analytical scale, following which the embryos were resuspended in a volume five times the mg weight of the sample of Cell Lysis Buffer (CLB; 10 mM HEPES, 10 mM NaCl, 1 mM KH_2_PO_4_, 5 mM NaHCO_3_, 4 mM EDTA, 1 mM CaCl_2_, 500 μM MgCl_2_, cOmpleteTM Protease inhibitor (Merck, 11836170001), phosphatase inhibitors (Merck, 4906845001), 10 μg/mL Granzyme B Inhibitor IV (Merck, 68056-1MG), pH 8.0). The embryos were then transferred, using a sterile glass pipette, into a Dounce Homogeniser (SLS, D8938-1SET), and the tissue homogenised with 15 turns of the A pestle. The tissue homogenate was then transferred into a fresh 1.5 mL tube and placed on ice for 15 minutes. To remove the yolk and debris, the samples were centrifuged at 300g for 5 minutes at 4°C. The supernatant was removed, and the cell pellet resuspended in a fresh 5x volume of CLB (as above). The cells were then transferred to the Dounce Homogeniser, and the cells were lysed with 15 strokes of the B pestle. The cell lysate was then transferred to a fresh 1.5mL tube on ice, and the total lysate fraction was made by adding a 2.5 mM Sucrose solution to the cell lysate at a 0.1 vol/vol ratio to the original CLB volume, mixing well. A portion of this was taken and stored at -80°C for further use as the total lysate. The samples were then centrifuged at 6,300g for 10 minutes at 4°C, with the supernatant collected and stored at -80°C for further use as the cytoplasmic fraction. The nuclear pellet was resuspended in 500 μL of Tris/Saline/EDTA/0.1% IGEPAL (TSE 0.1%; 8 mM Tris-HCl, 300 mM Sucrose, 1mM EDTA, 0.1% IGEPAL CA-630 (Merck-I8896)) buffer, and the samples centrifuged at 4,000g for 5 minutes at 4°C. The supernatant was removed, and the nuclear pellet washed three more times with 500μL TSE 0.1%. After the final wash, the nuclear pellet was resuspended in 500 μL of a TSE 1% IGEPAL buffer (TSE 1%), and placed on ice for 30 minutes with intermittent vortexing every 10 minutes. Following this, the nuclear lysate was sonicated using a Soniprep 150 plus Ultrasonic Disintegrator (MSE (UK) LTD, MSS150-CX4-5) at 23 kilohertz (kHz) for 15 seconds. The nuclear lysate was then stored at -80°C for further use.

### Western blotting

For Western blotting validation of the knock-in protein, 15 μg of total protein (per fraction) content was mixed into a solution containing 1X LDS Sample Buffer (Fisher Scientific, 13276499) and 1X Sample Reducing Agent (Fisher Scientific, 13296499). The samples were loaded onto a 4-12% Bis-Tris gel (Fisher Scientific, 10452342) alongside the PageRuler Prestained 10-250 kDa ladder (Fisher Scientific, 11832124). The gels were run at 160V for 10 minutes, followed by 180V for 50 minutes. The gels were then incubated in a 20% ethanol solution for 8 minutes, following which they were transferred to a PVDF membrane (Fisher Scientific, 15249296) using the iBlot2 transfer device (ThermoFisher, IB21001) and the default P0 programme. Membranes were then immunoassayed with the following primary antibodies: goat anti-GFP 1:2000 (Abcam, ab6673), rabbit anti-SNRNP70 1:500 (Sigma, AV40276), mouse anti-α tubulin 1:5000 (Cell Signalling, 3873T), and mouse anti-laminb1 1:1000 (Merck, 11243217001). For the detection of the primary antibodies, donkey anti-goat IgG (H+L) (Abcam, ab6885), goat anti-rabbit IgG (H+L) (Abcam, ab205718), and goat anti-mouse IgG (H+L) (Abcam, ab205719) HRP conjugated antibodies were used (all at 1:5000), and were detected using the PicoPlus Chemiluminescent Substrate (Fisher Scientific, 15626144). Membranes were visualised using the Bio-Rad ChemiDoc MP imaging system, using multiplex imaging with epifluorescence and chemiluminescence.

### Total RNA extraction

At 96 hpf, TgKI(*snrnp70-eGFP*) and *mitfa* embryos were anaesthetised, manually dechorionated, and 15 per biological replicate were transferred into a sterile 1.5 mL tube. The excess solution was removed, 350 μL of Buffer RLT from the RNeasy Micro kit (QIAGEN, 74004) was added, and the embryos homogenised using a sterile Bel-Art pestle (Merck, BAF199230001-100EA). Once the solution was clear, with no large debris, the homogenate was passed onto a QIAshredder homogenise column (79656) and centrifuged at maximum speed for 2 minutes at 4°C. The homogenised tissue was then taken through the RNEasy Micro Kit protocol. RNA concentration and purity was assessed using the RNA ScreenTape Kit (Agilent, 5067-5576) and the Agilent TapeStation 4200 device (Agilent, G2991BA).

### Reverse transcription quantitative PCR (RT-qPCR)

After total RNA isolation from TgKI(*snrnp70-eGFP*) and *mitfa* embryos, a first strand synthesis reaction was performed using Maxima First Strand Synthesis Kit (ThermoFisher, K1681), using 0.78 μg of input RNA per reaction. The RT-qPCR reaction was performed using the SYBR Green Master Mix (Fisher Scientific, A25743) on a StepOne Real Time PCR system (ThermoFisher, 4376357).

### Standard curves and Relative Quantification using the Pfaffl method

Standard curves for were generated for target and reference genes, using a five point five-fold serial dilution of first strand cDNA (with the first point being a 1/10 dilution of the neat cDNA). The Ct values were plotted against the negative logarithm of the template concentration, and percentage primer efficiency was calculated using the following equation:

Relative gene expression was calculated using the Pfaffl equation, which corrects for differences in amplification efficiency between the target and reference genes.

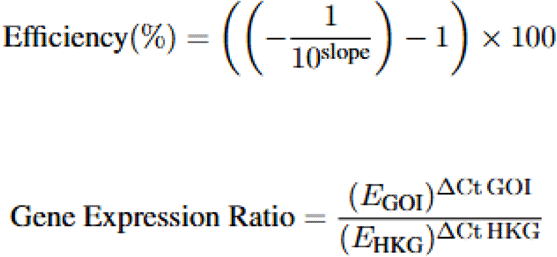

where ΔCt value was calculated by normalising against the housekeeping gene, rps29, using the equation ΔCt = Average Control Ct - Average target Ct.

### Confocal microscopy

#### Imaging of founders

Imaging of F_1_ and F_2_ TgKI(*snrnp70-eGFP*) embryos was carried out using an Olympus BX63LF/FV3000 upright confocal microscope equipped with a XLUMPLFLN20XW 20x/1.0 NA water immersion objective (Olympus). Excitation was provided at 488nm for enhanced green fluorescent protein (eGFP). High resolution images were taken 0.31 x 0.31 μm resolution (1024 x 1024 pixels).

#### Imaging of IHC-IF embryos

Imaging of IHC-IF cryosections on F_2_ SNRNP70eGFP embryos was carried out using the Olympus BX63LF/FV3000 upright confocal microscope equipped with a UPLSAPO 40x/0.95 air objective (Olympus). Excitation was provided using 461nm (for DAPI), 520nm (for Alexa-Fluor 488), 603 nm (for Alexa-Fluor 568), and 650 nm (for Alexa-Fluor 633). High resolution images were taken at 0.31 x 0.31 μm resolution (1024 x 1024 pixels).

#### Imaging of ICC-IF 12DIV cultures

Imaging of ICC-IF on SNRNP70eGFP primary neuronal cultures was carried out using the Olympus BX63LF/FV3000 upright confocal microscope equipped with a PLN100XO 100x/1.25 Oil objective (Olympus). Excitation was provided using 461nm (for DAPI), 520nm (for Alexa-Fluor 488), 603 nm (for Alexa-Fluor 568), and 650 nm (for Alexa-Fluor 633). High resolution images were taken at 0.1 x 0.1 μm resolution (1024 x 1024 pixels).

#### Imaging of PLA 12DIV cultures

Imaging of PLA on SNRNP70eGFP primary neuronal cultures was carried out using the Olympus BX63LF/FV3000 upright confocal microscope equipped with a PLN100XO 100x/1.25 Oil objective (Olympus). Excitation was provided using 461nm (for DAPI), 520nm (for Alexa-Fluor 488), 603 nm (for Alexa-Fluor 456), and 650 nm (for Alexa-Fluor 633). High resolution images were taken at 0.1 x 0.1 μm resolution (1024 x 1024 pixels).

#### Time-lapse imaging of SNRNP70 within RNP granules

Fertilised TgKI(*snrnp70-eGFP*) or TgKI(*snrnp70-eGFP*);Tg(*mnx1:mCherry*) embryos at 1-cell stage were injected with a 100 μM Cy5-UTP solution. Embryos at 48 hpf were mounted in 1% low melting point agarose (ThermoFisher Scientific, 16520050). Imaging was performed using a Zeiss LSM 880 Fast Airyscan confocal microscope equipped with GaAsP spectral detectors and a 40x/1.2 N.A. water-immersion objective (Carl Zeiss). Excitation was provided using 488 nm (for GFP), 561 (for mCherry) and 633 nm (for Cy5) solid state lasers. High resolution images were captured at 2.6 Hz at a 0.04 x 0.04 μm resolution (488 x 488 pixels) before being processed for Airyscan imaging.

### Data analysis

#### PLA quantification

Images were separated by channel and z-stack prior to processing in CellProfiler (Stirling et al., 2021). Nuclear regions were first identified using the ‘**IdentifyPrimaryObjects’** module with the DAPI channel. Subsequently, neuronal somata were defined using the ‘**IdentifySecondaryObjects’** module, with the acetylated α-Tubulin (AcTub) channel used as the identifying image and nuclei used as the primary objects. The soma cytoplasmic region was then delineated using the ‘**IdentifyTertiaryObjects’** module by defining the soma as the larger object and the nucleus as the smaller object.

To identify axonal compartments, the Ac α-Tub channel was first used with the ‘**IdentifyPrimaryObjects’** module to define Ac α-Tub-positive structures as primary objects. Axonal regions were subsequently separated from soma using the ‘**IdentifyTertiaryObjects’** module by defining Ac α-Tub-positive structures as the larger object and the soma as the smaller object.

Following segmentation of cellular compartments, the PLA channel was enhanced using the ‘**EnhanceOrSuppressFeatures’** module to improve detection of PLA puncta. The enhanced PLA image was then analysed using the ‘**IdentifyPrimaryObjects’** module to identify individual PLA puncta. PLA puncta were subsequently quantified within the three defined cellular compartments: nucleus, soma cytoplasm, and axons.

To normalise PLA signal between compartments, the surface area of each compartment was calculated using the ‘**MeasureImageAreaOccupied’** module. Pixel counts were converted to μm² based on the image calibration settings. The total number of PLA puncta and the corresponding compartment area were summed across all z-planes, and PLA signal was expressed as the number of PLA puncta per μm² for each cellular compartment.

#### Statistical analysis

Definition of statistical significance is indicated in the figure legends. For PLA quantifications, one-sided t-test was used to compare differences between experimental and control (anti-GFP antibody alone) groups. For comparing gene expression ratios, Welch’s t-test was used. For quantification of the percentage number of eGFP+ puncta co-localising with the mRNP granule marker, a two-tailed unpaired Mann-Whitney test was used. The criterion for statistical significance was set at *p* < 0.05 and results are represented as mean ± SEM.

## Results

### Generation and validation of an endogenous SNRNP70-eGFP knock-in zebrafish line

To facilitate visualisation of the endogenous SNRNP70 protein and characterise its expression and localisation *in vivo* and *in vitro*, we generated a CRISPR-mediated knock-in (KI) zebrafish line in which an eGFP sequence was fused to the C-terminus of the endogenous *snrnp70* gene, following the endogenous tagging approach described by Levic et al. (2021) (Levic et al., 2021) (Fig.1A). Briefly, the donor construct was designed to introduce a C-terminal eGFP tag in-frame with the SNRNP70 protein, as well as preserving the endogenous regulatory elements of the gene (i.e., by containing the endogenous 3’UTR as a 3’ homology arm). Founder animals were identified based on eGFP fluorescence in the F_1_ offspring, indicating successful germline transmission. Sequencing analysis of the KI allele confirmed the presence of *egfp* sequence between exon 10 (last exon of *snrnp70*) and 3’UTR indicating successful integration (Fig.S1). The sequencing also revealed a small insertion-deletion mutation (INDEL) downstream of the integration boundary, however this alteration was within a non-coding region (intron 9) (Fig.S1).

**Figure 1:**
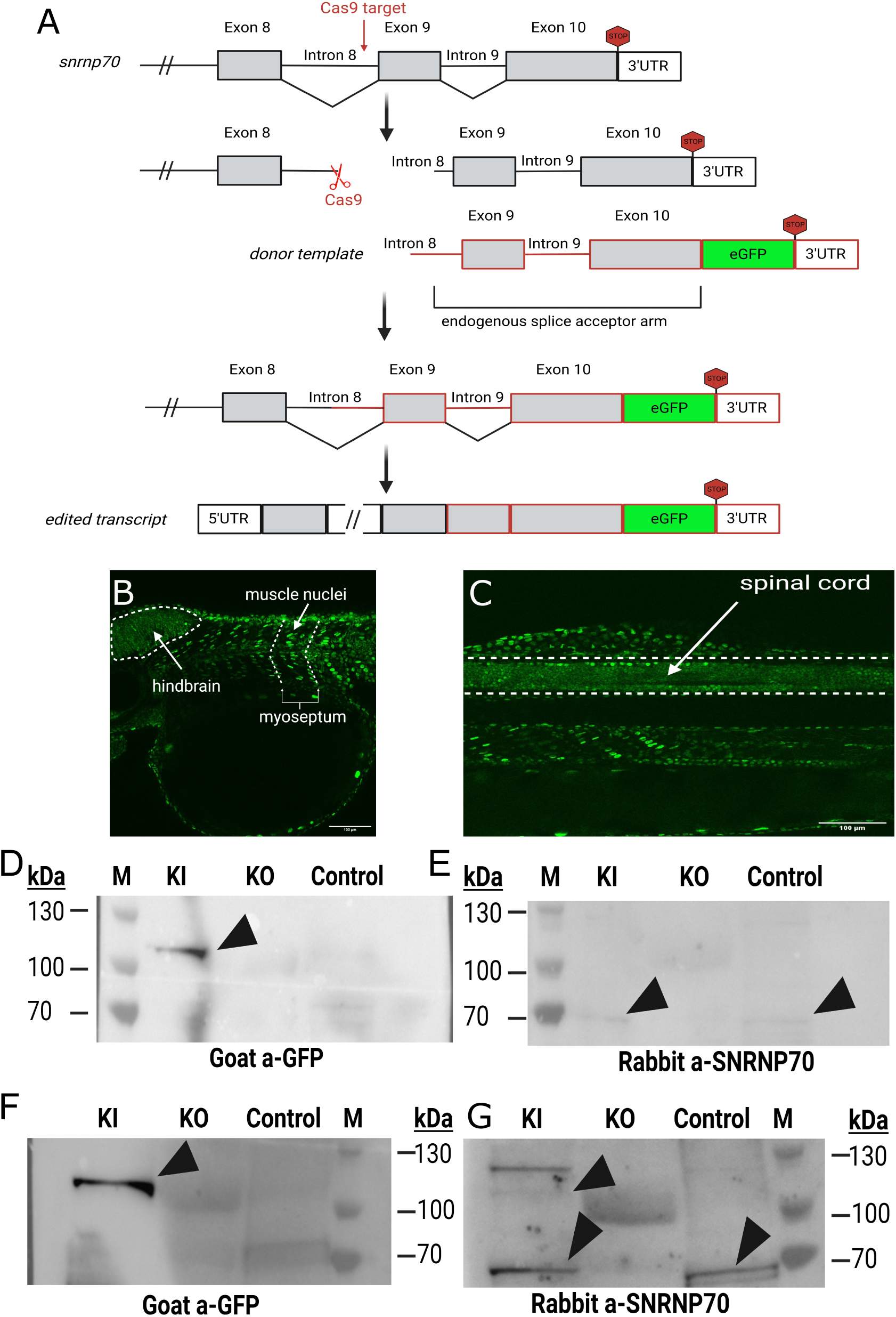
Generation and validation of an endogenous SNRNP70-eGFP knock-in zebrafish line. **(A)** Genomic architecture of the *Danio rerio snrnp70* locus and schematic diagram of the C-terminal eGFP tagging strategy. The gRNA/Cas9 nuclease target is located 80bp upstream of the 3’ end of intron 9. The best available intron, based on design constraints from Levic et al. (2021), was targeted to induce a double stranded DNA (dsDNA) break, following which a donor template containing a 5’ splice acceptor element is integrated (likely through NHEJ). Integration of the cassette and mRNA splicing results in the expression of the eGFP tagged protein. **(B)** Lateral image of TgKI(*snrnp70-eGFP*) heterozygous F_1_ larva at 48 hpf. Expression of the SNRNP70-eGFP protein is observed throughout the hindbrain (left arrow and outline), along with muscle nuclei in the myosepta (right arrow). Scale bar, 100 μm. **(C)** Lateral image of TgKI(*snrnp70-eGFP*) heterozygous F_1_ larva at 48 hpf. Expression of the SNRNP70-eGFP protein seen throughout the spinal cord. Scale bar, 100 μm. **(D)** Detection of the eGFP-tagged SNRNP70 protein in the total lysate by Western blot. Total lysate samples from knock-in (KI), knock-out (KO) and mitfa (control) embryos were analysed using a goat anti-GFP antibody. A specific immunoreactive band was observable at ∼110kDa in the KI sample, as indicated by the left arrowhead. No corresponding band was observed in the KO or control samples. Lane identities are indicated above the blot. **(E)** Detection of the native SNRNP70 protein in the total lysate, visualised on a rabbit anti-SNRNP70 blot. Specific immunoreactive bands at ∼70kDa are visible in both the KI and control lanes (left and right arrowheads), and not visible within the KO lane. **(F)** Immunoblot using a goat anti-GFP antibody showing the detection of a GFP immunoreactive band at ∼110kDa in the cytoplasmic fraction (left arrowhead). **(G)** Detection of both the SNRNP70-eGFP fusion protein and the native SNRNP70 proteins within the cytoplasmic fraction. Upper left arrow shows a specific immunoreactive band at ∼110kDa, indicative of the fusion protein. The lower banks in both the KI and control lanes indicate the native SNRNP70 proteins.

F_1_ TgKI(*snrnp70-eGFP*) progeny exhibited robust eGFP fluorescence throughout the axis of the embryo, consistent with expression of the integrated SNRNP70-eGFP allele and the established expression pattern of endogenous SNRNP70 (Nikolaou et al., 2022; Thomas-Jinu et al., 2017). Fluorescence was readily detectable throughout the developing brain and spinal cord, as well as across muscle fibres, demonstrating widespread expression of the tagged protein (Fig.1B and 1C). At the subcellular level, SNRNP70-eGFP fluorescence was particularly prominent within nuclei (Fig.1B), consistent with the established role of SNRNP70 as a core component of the spliceosome and its major nuclear function in pre-mRNA splicing. Together, these expression patterns provide an initial indication that the KI line faithfully recapitulates the expected broad and predominantly nuclear distribution of endogenous SNRNP70 in developing zebrafish embryos.

To determine whether insertion of the eGFP tag affected relative allele expression, we compared the expression levels of native (Exon10/3′UTR) and KI (Exon10/eGFP) transcripts by qPCR in heterozygous TgKI(*snrnp70-eGFP*) animals. We observed no evidence of a difference in expression between the eGFP-tagged and native transcripts (Welch’s t-test; t(3.53) = 0.716, p = 0.519; mean Exon10/3′UTR ratio = 1.046, mean Exon10/eGFP ratio = 0.940; 95% CI of the difference: −0.325 to 0.536; Fig.S2A). These findings indicate that the eGFP-tagged allele is expressed at levels similar to the native allele, supporting the use of the SNRNP70-eGFP allele as an endogenous reporter without evidence that insertion of the tag substantially alters *snrnp70* transcript abundance.

Expression of *lin7b*, the nearest downstream neighbouring gene, was similar between TgKI(*snrnp70-eGFP*) and *mitfa* control animals (Welch’s t-test; t(3.57) = −0.79, p = 0.480; mean TgKI(*snrnp70-eGFP*) = 0.82 ± 0.28 SEM; mean *mitfa* = 1.05 ± 0.40 SEM; Fig.S2B). These findings indicate that insertion of the eGFP KI cassette does not detectably alter expression of the neighbouring *lin7b* gene, supporting the conclusion that the KI does not cause a broader disruption of local gene expression at the *snrnp70* locus.

To confirm expression and subcellular localisation of the eGFP-tagged SNRNP70 protein, lysates from heterozygous TgKI(*snrnp70-eGFP*) embryos were analysed by Western blotting. Immunoblotting of total protein lysates with an anti-GFP antibody detected a single band at approximately 110 kDa in TgKI(*snrnp70-eGFP*) embryos, consistent with the predicted molecular weight of SNRNP70-eGFP (Fig.1D, lane 1). This band was absent from both *snrnp70* knockout and *mitfa* control embryos (Fig.1D, lanes 2 and 3, respectively), confirming the specificity of the GFP signal. Immunoblotting with an SNRNP70-specific antibody detected a single band of approximately 70 kDa in both *mitfa* control and TgKI(*snrnp70-eGFP*) embryos (Fig.1E, lanes 1 and 3), corresponding to the predicted molecular weight of native SNRNP70. This band was absent in *snrnp70* knockout embryos (Fig.1E, lane 2), further confirming antibody specificity and the identity of the detected protein.

Given previous evidence that SNRNP70 is also present in extranuclear neuronal compartments (Nikolaou et al., 2022; Thomas-Jinu et al., 2017), we next performed subcellular fractionation to determine whether the tagged protein was detectable within the cytoplasmic compartment. Western blot analysis of the cytoplasmic fraction revealed both eGFP-tagged and native SNRNP70 in heterozygous TgKI(*snrnp70-eGFP*) embryos (Fig.1F and 1G, lane 1), whereas the *mitfa* control contained only native SNRNP70 (Fig.1G, lane 3). Together, these findings confirm expression of the SNRNP70-eGFP fusion protein in TgKI(*snrnp70-eGFP*) embryos and demonstrate that the tagged protein is detectable in the cytoplasmic compartment in addition to its predominant nuclear localisation. Moreover, these results provide biochemical validation of the SNRNP70-eGFP KI line, supporting its use as an endogenous reporter of SNRNP70 localisation.

### Endogenous SNRNP70-eGFP is enriched in neuronal processes

Having established and validated the TgKI(*snrnp70-eGFP*) line, we next used this resource to define the spatial distribution of endogenous SNRNP70 in the developing nervous system. We assessed SNRNP70-eGFP distribution at 96 hpf using immunohistochemistry/immunofluorescence.

Within sections taken at the level of the forebrain, SNRNP70-eGFP protein immunoreactivity was detectable across all diencephalic neurons (Fig.2A). Intriguingly, eGFP fluorescence was also observed within the retina, with strong signal visible in the outer nuclear layer (ONL; containing the rod and cone photoreceptor cell bodies) and the inner plexiform layer (IPL; containing the synaptic connections between the bipolar cell axons, amacrine neurites and retinal ganglion cell dendrites) (Fig.2A). Fluorescence was also visible in the outer plexiform layer (OPL; containing the synaptic connections between the bipolar cells, horizontal cells and photoreceptors) and the retinal ganglion cell layer (GCL; harbouring the somata of retinal ganglion cells) (Fig.2A).

**Figure 2:**
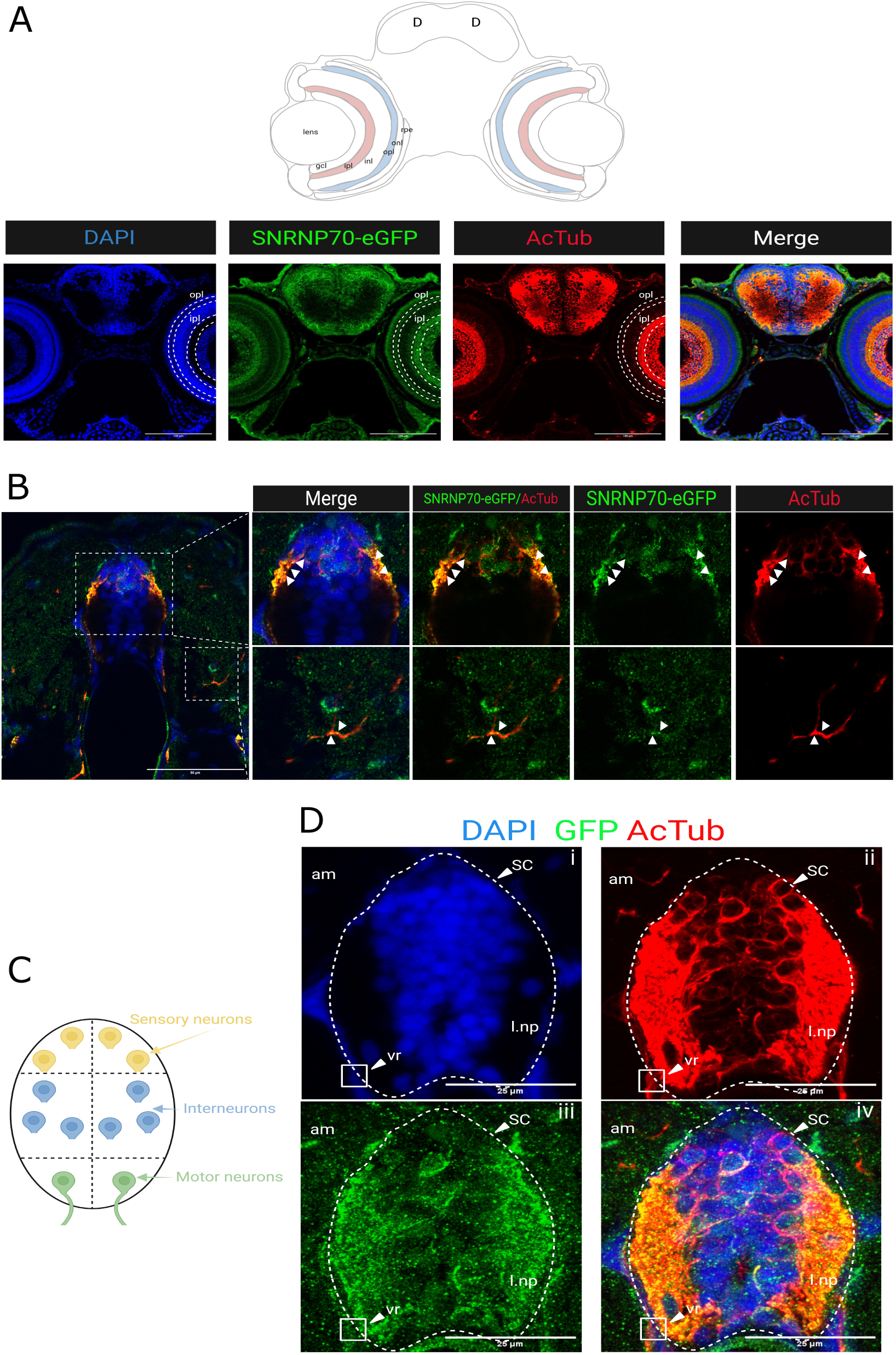
Endogenous SNRNP70-eGFP is enriched in neuronal processes. **(A)** IHC-IF staining for SNRNP70-eGFP, showing widespread distribution throughout the forebrain and retina. SNRNP70-eGFP is particularly expressed in the outer plexiform layer (opl) and inner plexiform layer (ipl) of the retina. Schematic representation adapted from https://zebrafishucl.org/retina. From left to right: DAPI, acetylated α-Tubulin, SNRNP70-eGFP and merge of all channels. D, diencephalon; gcl, ganglion cell layer; inl, inner nuclear layer; ipl, inner plexiform layer; onl, outer nuclear layer; opl, outer plexiform layer; rpe, retinal pigment epithelium. Scale bar, 100 μm. **(B)** IHC-IF images showing the localisation of SNRNP70-eGFP protein in the dorsal spinal cord. Upper inset is within the dorsal spinal cord. From left to right: Merge of all channels/DAPI, acetylated α-tubulin/SNRNP70-eGFP, SNRNP70-eGFP, and acetylated α-tubulin. Arrowheads indicate fluorescent puncta containing SNRNP70-eGFP within the lateral synaptic regions of the spinal cord. Lower inset shows motor neuron axon branches in the area of the horizonal myoseptum. Arrows point towards SNRNP70-eGFP fluorescent puncta within motor neuron processes (positive for acetylated α-Tubulin). Scale bar, 50 μm. **(C)** Schematic diagram depicting an outline of the organisation of the spinal cord at 96 hpf. The location of sensory neurons, interneurons and motor neurons within the spinal cord is shown. **(D)** IHC-IF images showing SNRNP70-eGFP fluorescence within spinal cord neuronal somata, lateral neuropil and ventral root area. (i) DAPI counterstain, (ii) acetylated α-Tubulin, (iii) SNRNP70-eGFP, (iv) merge of all signals. Axial muscle (am); lateral neuropil (l.np); spinal cord (sc); ventral root (vr). Scale bar, 25 μm.

At the level of the trunk, SNRNP70-eGFP was observed both within the dorsal spinal cord as well as enriched within the lateral synaptic regions of the spinal cord, containing predominantly neuronal processes (Fig.2B, upper inset and panels). Punctate SNRNP70-eGFP^+^ formations could be observed within these neuronal processes marked by acetylated α-Tubulin (Fig.2B, upper panels, left and right arrows). Moreover, a punctate SNRNP70-eGFP fluorescence was also observed within peripheral motor neuron projections in the medial myotome (Fig.2B, lower panels).

Higher-magnification images of the spinal cord revealed somatic GFP fluorescence throughout the dorso-ventral extent of the neural tube (Fig.2Di and 2Diii), indicating the presence of SNRNP70-eGFP in sensory neurons, interneurons and motor neurons (Fig.2Di). Surprisingly, strong expression of the SNRNP70-eGFP protein could be seen within the lateral neuropil (Fig.2Diii), as well as within the ventral root region (Fig.2Dii and 2Div, lower left arrow).

Together, these analyses demonstrated widespread expression of endogenous SNRNP70-eGFP throughout the developing zebrafish nervous system. Although strong nuclear fluorescence was observed across neuronal populations, SNRNP70-eGFP was also prominent within neuronal processes, including synaptic neuropil regions in the retina and spinal cord, ventral roots and peripheral motor neuron projections. The presence of discrete punctate SNRNP70-eGFP^+^ structures within these processes was particularly notable, raising the possibility that endogenous SNRNP70 associates with RNA-containing complexes outside the nucleus.

### Endogenous SNRNP70 localises to dynamic neuronal mRNP granules

The punctate distribution of SNRNP70-eGFP suggested that endogenous SNRNP70 may associate with neuronal mRNP granules. Previous work demonstrated that transgenically expressed human SNRNP70 localises to mRNP granules and co-localises with mRNA targets in neurons (Nikolaou et al., 2022). The endogenous KI reporter therefore provides an opportunity to investigate this association under physiological protein abundance and in the native genomic context. We used the TgKI(*snrnp70-eGFP*) line to visualise endogenous SNRNP70 *in vivo*. mRNP granules were labelled by microinjection of a Cy5-conjugated uridine-5′-triphosphate analogue (Cy5-UTP), which is incorporated into newly synthesised RNA and subsequently labels cytoplasmic mRNP granules (Nikolaou et al., 2022; Wong et al., 2017). At 48 hpf, SNRNP70-eGFP showed strong nuclear localisation (asterisks in Fig.3A), together with numerous extranuclear eGFP^+^ puncta throughout the posterior brain and spinal cord. These punctate structures likely represent the endogenous cytoplasmic pool of SNRNP70 and provided an opportunity to directly assess its association with mRNP granules under physiological expression conditions.

**Figure 3:**
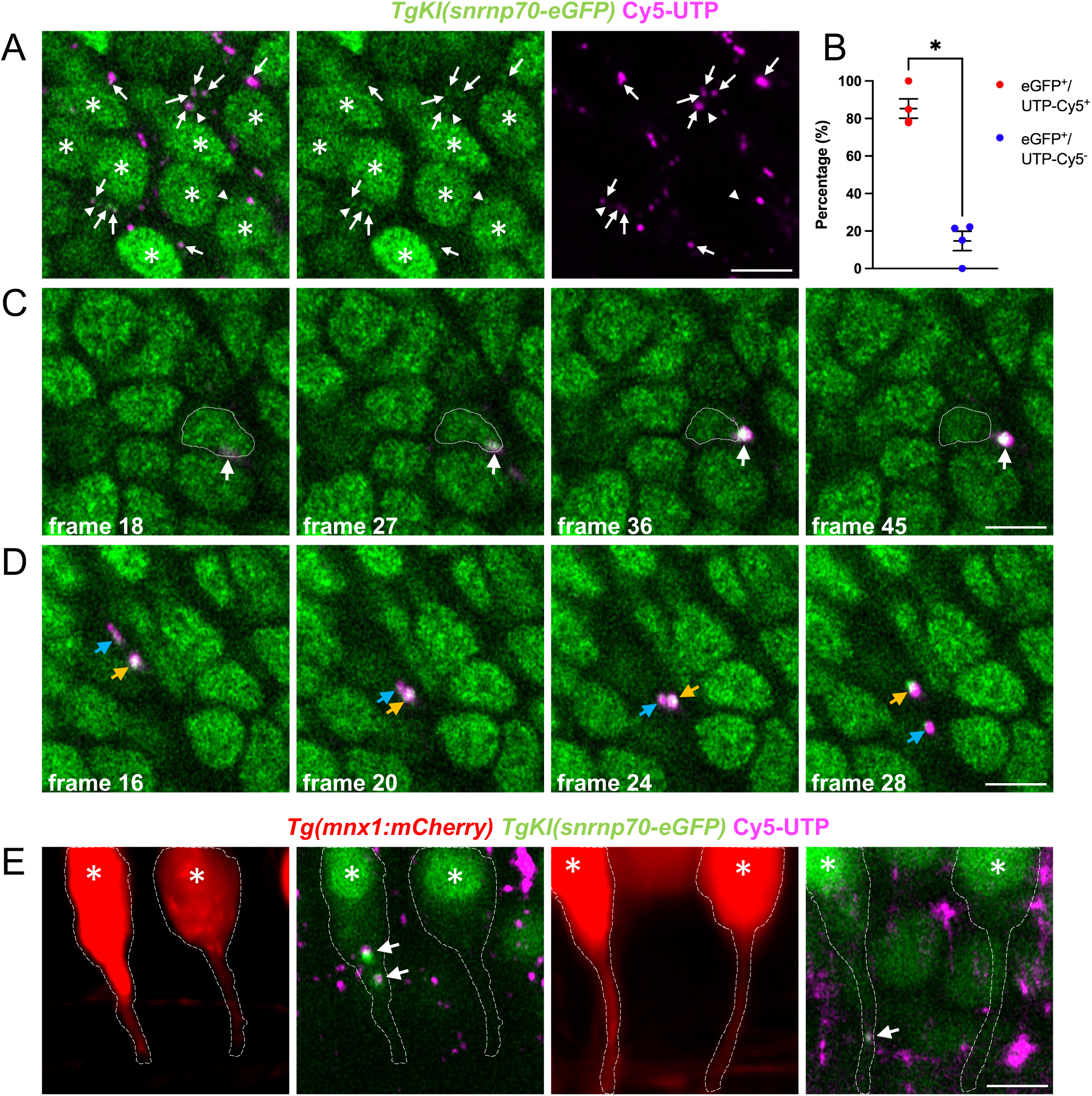
Endogenous SNRNP70 localises to dynamic neuronal mRNP granules. **(A)** Representative example of TgKI(*snrnp70-eGFP*) showing the live *in vivo* localisation of SNRNP70-eGFP in the hindbrain region at 48 hpf. Cy5-UTP marks mRNP granules. Asterisks indicate eGFP fluorescence within nuclei. Arrows point to puncta that are positive for both eGFP and Cy5. Arrowheads indicate eGFP^+^ puncta alone. Scale bar, 5 μm. **(B)** Quantification showing the percentage number of eGFP^+^ puncta found alone or co-localising with the mRNP granule marker. Graph shows mean values ± SEM. *p < 0.05, two-tailed unpaired Mann-Whitney test. n = 4 animals in two independent experiments. **(C)** Time-lapse of TgKI(*snrnp70-eGFP*) hindbrain neurons at 48 hpf. Representative side view images showing the appearance of a GFP^+^ mRNP granule (arrow). Dashed line demarcates the boundaries of the nucleus. Scale bar, 5 μm. **(D)** Time-lapse of TgKI(*snrnp70-eGFP*) hindbrain neurons at 48 hpf. Representative side view images showing dynamic GFP^+^ mRNP granules (arrows). Scale bar, 5 μm. **(E)** Representative side view images of TgKI(*snrnp70-eGFP*);Tg(*mnx1:mCherry*) larvae at 48 hpf depicting the localisation of SNRNP70-eGFP within motor neurons in the ventral spinal cord. eGFP^+^ mRNP granules (arrows) can be seen in both the soma (asterisk depicting the nucleus) and proximal axon of motor neurons. Scale bar, 5 μm.

We next asked whether the extranuclear SNRNP70-eGFP puncta represented mRNP granules. SNRNP70-eGFP puncta frequently overlapped with Cy5-labelled mRNP granules (Fig.3A), and quantitative analysis confirmed that the majority of extranuclear SNRNP70-eGFP^+^ puncta were associated with Cy5-labelled mRNP granules (Fig.3B), providing evidence that endogenous SNRNP70 is preferentially localised to RNA-rich granules *in vivo*. Time-lapse imaging further revealed that SNRNP70-eGFP^+^ mRNP granules were highly dynamic (Video S1 and S2), displaying both short-range movements within the perinuclear region (Fig.3C) and sustained long-range transport (Fig.3D). These observations demonstrate that endogenous SNRNP70 is associated with dynamic and motile mRNP granules, supporting a role for SNRNP70 in the organisation and intracellular trafficking of RNA-containing complexes.

Crossing the endogenous SNRNP70 reporter with a motor-neuron-specific mCherry reporter further demonstrates the compatibility of the KI line with cell-type-specific genetic tools. To this end, we crossed the TgKI(*snrnp70-eGFP*) line with the Tg(*mnx1:mCherry*) motor neuron reporter line. Imaging revealed numerous Cy5-labelled mRNP granules within motor neuron somata and proximal axons, with a substantial proportion also displaying SNRNP70-eGFP fluorescence (arrows in Fig.3E). Together, these findings extend previous observations based on transgenic SNRNP70 expression by demonstrating that SNRNP70-eGFP from the endogenous knock-in allele is readily detected within dynamic neuronal mRNP granules *in vivo*. Importantly, the KI approach demonstrates that this localisation is maintained under endogenous expression conditions, providing strong evidence that the association of SNRNP70 with mRNP granules is a physiological property of the protein rather than a consequence of overexpression.

### Endogenous SNRNP70 is in close proximity to mRNP granule-associated RBPs in neuronal compartments

Given the punctate distribution of SNRNP70-eGFP within neuronal processes and its localisation to mRNP granules, we next asked whether endogenous SNRNP70 co-localises and associates with established components of neuronal mRNP granules. We focused on PABPC1B, FUS, and UPF1, which have all been implicated as components of neuronal mRNP granules (Ravanidis et al., 2018). To assess the distribution and relationship of these RBPs with SNRNP70, we performed immunofluorescence analysis using commercially available antibodies against each protein. The selected antibodies are either predicted to recognise the corresponding zebrafish orthologues (FUS and UPF1) or were raised against immunogens derived from zebrafish protein sequences (PABPC1B), enabling assessment of their localisation and association with endogenous SNRNP70 in the zebrafish nervous system.

As expected, immunostaining of primary zebrafish embryonic neurons revealed distinct subcellular distributions among the mRNP granule-associated proteins. PABPC1B and UPF1 were enriched throughout the soma and axons (Fig.S3A and S3C, respectively). In contrast, FUS was predominantly nuclear, with diffuse staining extending into axons (Fig.S3B). Consistent with these established localisation patterns, immunohistochemical analysis of 96 hpf TgKI(*snrnp70-eGFP*) larvae revealed extensive co-localisation of endogenous SNRNP70 with FUS, PABPC1B, and UPF1 within discrete punctate cytoplasmic structures (Fig.S4-S6). These findings further support the association of endogenous SNRNP70 with neuronal mRNP granules and indicate that SNRNP70 occupies structures containing multiple established components of the mRNP granule machinery.

To determine whether endogenous SNRNP70 is in close molecular proximity to established neuronal mRNP granule components, we performed proximity ligation assays (PLA) against PABPC1B, FUS, and UPF1. Primary zebrafish embryonic neurons derived from TgKI(*snrnp70-eGFP*) embryos enabled robust segmentation of distinct cellular compartments - the nucleus, soma cytoplasm and neurites - and quantitative analysis of PLA signal within each compartment. Moreover, by preserving endogenous expression and localisation, this novel resource avoids potential artefacts associated with overexpression while providing an independent reporter to complement antibody-based approaches, which can be limited by antibody specificity and sensitivity. PLA puncta, indicative of close proximity between SNRNP70 and PABPC1B (Fig. 4A), FUS (Fig. 4B), and UPF1 (Fig. 4C), were significantly enriched in the nucleus, somatic cytoplasm, and neurites compared with the GFP-antibody-only control. These findings suggest that SNRNP70 is in close proximity to these RBPs across multiple neuronal compartments. Together, these findings identify RBPs as candidate proteins associated with endogenous SNRNP70 and demonstrate the utility of the KI line for investigating SNRNP70-protein proximity across multiple neuronal compartments. More broadly, these experiments demonstrate the utility of the SNRNP70-eGFP KI line as a physiologically relevant model for investigating the molecular environment of endogenous SNRNP70. Together with the live-imaging capabilities of eGFP, this resource provides a versatile platform for investigating the localisation, dynamics and molecular associations of endogenous SNRNP70 in neuronal mRNP granules.

**Figure 4:**
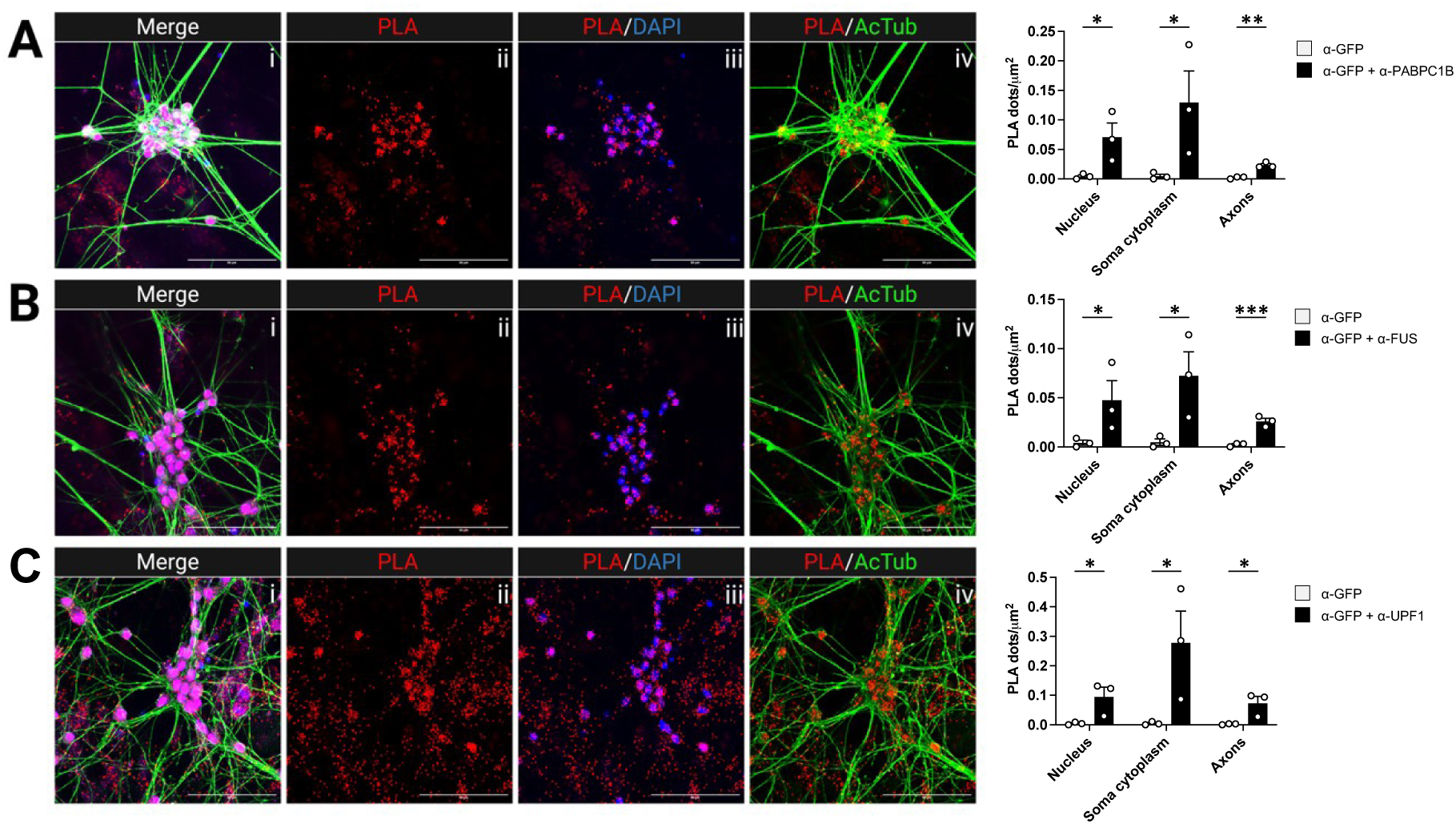
Endogenous SNRNP70 is in close proximity to mRNP granule-associated RBPs in neuronal compartments. Representative proximity ligation assay (PLA) images of 12 DIV cultured primary neurons derived from TgKI(*snrnp70-eGFP*) zebrafish embryos indicating the interaction between **(A)** SNRNP70 and PABPC1B; **(B)** SNRNP70 and FUS; and **(C)** SNRNP70 and UPF1. PLA puncta (red) indicate sites of protein proximity, acetylated α-Tubulin (green) labels neuronal processes, and DAPI (blue) stains nuclei. The graphs on the right-hand side show quantification of PLA signals between SNRNP70-GFP and each of the candidate interacting protein. Bar graphs show the mean PLA puncta density (PLA dots/µm² ± SEM) in the nucleus, soma cytoplasm and neurites. Statistical significance was assessed using one-sided t-test. Asterisks indicate statistically significant differences: * *P* < 0.05; ** *P* < 0.01; *** *P* < 0.001. Scale bar, 50 μm.

## Discussion

Here, we describe the generation and validation of a CRISPR-mediated zebrafish KI line in which eGFP was fused to the C-terminus of endogenous SNRNP70, enabling fluorescent visualisation of endogenous SNRNP70 in living neurons. By preserving the native 3′UTR and endogenous regulatory context, this TgKI(*snrnp70-eGFP*) line enables visualisation of SNRNP70 under conditions that more closely reflect its physiological expression, localisation, function, and regulation (Boswell et al., 2025; Levic et al., 2021). This is particularly important for the study of granule-associated RBPs, whose localisation and dynamics can be highly sensitive to protein abundance and cellular context, and for which overexpression may itself promote aberrant granule formation (Maharana et al., 2018; Pushpalatha et al., 2022; Sanders et al., 2020). The KI strategy therefore provides an alternative approach to conventional visualisation methods, enabling SNRNP70 to be monitored at endogenous expression levels within its native genomic context while avoiding artefacts associated with overexpression and variability arising from transgene integration site and copy number (Roberts et al., 2014).

Despite these advantages, CRISPR-mediated KI approaches can be technically challenging and may introduce unwanted INDELs during the integration process. Importantly, such INDELs were also observed at the integration boundaries in the approach described by Levic et al. (2021), who demonstrated that these alterations can be tolerated when they occur within sequences that are subsequently removed during RNA splicing. Consistent with this, we detected a small INDEL downstream of the integration boundary in our TgKI(snrnp70-eGFP) line. However, this sequence alteration had no detectable effect on SNRNP70 protein expression levels, likely because the INDEL affected a non-coding region that is removed during splicing.

An important feature of the SNRNP70-eGFP line is its utility extends beyond fluorescence-based visualisation, and shows excellent compatibility with complementary genetic, imaging and biochemical approaches. Endogenous SNRNP70-eGFP provided a defined source of biomaterial for downstream analyses and enabled immunofluorescence detection using antibodies directed against the eGFP tag, providing a complementary and readily standardised approach for visualising endogenous SNRNP70 across experimental contexts. Moreover, the KI line was compatible with several orthogonal approaches, including Cy5-UTP microinjection to label nascent RNA and visualise mRNP granule dynamics *in vivo* (Nikolaou et al., 2022; Wong et al., 2017), genetic crossing with the Tg(*mnx1:mCherry*) motor-neuron reporter to assign cell-type identity to SNRNP70-eGFP^+^ granules, and derivation of primary neuronal cultures for proximity ligation assay (PLA)-based interaction studies. Together, these approaches establish a multimodal experimental pipeline that combines endogenous protein tagging, live imaging of RNA-containing granules, cell-type-specific reporter analysis, immunofluorescence, and culture-based protein-interaction assays. Importantly, the convergence of these independent approaches strengthens the interpretation that the observed localisation and interactions reflect properties of endogenous SNRNP70 rather than consequences of ectopic expression.

Using this KI line, we demonstrate that endogenous SNRNP70 is broadly expressed throughout the nervous system, but displays marked regional enrichment and a prominent punctate or granular distribution within neuronal processes. Rather than being predominantly nuclear, SNRNP70 was readily detected in axonal and synaptic regions such as the inner and outer plexiform layers of the retina, the lateral neuropil of the spinal cord, motor-neuron projections, and the ventral roots. The enrichment of SNRNP70 within these regions is particularly notable given that the inner plexiform layer and spinal neuropil comprise dense networks of dendritic and synaptic processes (Randlett et al., 2013), while motor-axon projections and terminals are established sites of local RNA regulation and translation (Briese et al., 2020; Tu et al., 2024). Together, these findings provide anatomical evidence supporting a physiological cytoplasmic role for SNRNP70 in neuronal post-transcriptional RNA regulation, extending beyond its established nuclear functions (Soares et al., 2025).

Utilising the novel TgKI(*snrnp70-eGFP*) line, we demonstrate that endogenous SNRNP70 is readily detected within neuronal Cy5⁺ RNA-containing mRNP granules *in vivo*. These SNRNP70-eGFP^+^ granules exhibited heterogeneous dynamics, with some undergoing shorter-range movements in perinuclear regions and others displaying longer-range trafficking throughout the neuronal cytoplasm and neurites. Complementary findings from our laboratory (Baldacchino et al., 2026), using the same endogenous SNRNP70-eGFP resource, independently demonstrate localisation of SNRNP70 within mRNP granules. Together, these studies illustrate the utility of the knock-in line for resolving the localisation and dynamics of endogenous SNRNP70 in neuronal RNA-containing complexes, and support that this observation is not dependent on ectopic SNRNP70 expression. This further substantiates previous observations from an overexpression model showing that SNRNP70 localises to motile mRNP granules in zebrafish motor axons and may contribute to the local regulation of transcript stability and processing (Nikolaou et al., 2022), while demonstrating that this localisation is maintained under endogenous expression conditions.

The association of endogenous SNRNP70 with dynamic RNA-containing granules, together with its proximity to PABPC1B, FUS, and UPF1, supports its presence within neuronal mRNP granule-like structures, established sites of mRNA transport, storage, and translational regulation (Bauer et al., 2023; De Graeve & Bessé, 2018; Formicola et al., 2019). Together with the presence of SNRNP70 in synaptic, dendritic, and axonal compartments of retinal and spinal cord neurons, these observations suggest that extranuclear SNRNP70 may contribute to neuronal mRNP granule biology and spatially regulated post-transcriptional RNA control, including RNA transport, localisation, stability, and local processing.

The proximity of SNRNP70 to PABPC1B and FUS suggests a potential role in the dynamic regulation of mRNA localisation, stability, and translation. PABPC1B promotes mRNA stability and translation through poly(A)-tail binding and interaction with eIF4G (Safaee et al., 2012), while FUS binds neuronal mRNAs, often recognising G-quadruplex structures, and facilitates their transport in mRNP complexes along neurites (Deshpande et al., 2019; Imperatore et al., 2020). FUS also forms cytoplasmic RNA granules that recruit mRNA complexes and support local protein synthesis (Yasuda et al., 2013). The association of SNRNP70 with these proteins therefore raises the possibility that it is incorporated into mRNP transport granules, where it may contribute to transcript storage and protection. Consistent with this model, SNRNP70 has been shown to confer transcript-selective protection from degradation (Nikolaou et al., 2022), suggesting that its RNA-regulatory functions may extend beyond splicing to include spatial control of transcript stability and translation.

The observed proximity between SNRNP70 and UPF1 raises an interesting hypothesis for future investigation. Given the established and central roles of UPF1 in nonsense-mediated mRNA decay (NMD), a major cellular RNA surveillance pathway (He & Jacobson, 2015; Hug et al., 2015), together with NMD-independent RNA-regulatory functions in mRNA transport, localisation, and local translation (Graber et al., 2017), SNRNP70 may potentially influence UPF1-associated mRNA fate pathways. Such a mechanism could provide a potential explanation for the transcript-selective protection from degradation previously attributed to SNRNP70 (Nikolaou et al., 2022). Determining whether SNRNP70 directly regulates UPF1 activity, recruitment or substrate selection will require additional functional and biochemical experiments.

Overall, this study establishes a validated endogenous SNRNP70-eGFP zebrafish line as a versatile resource for studying SNRNP70 localisation, dynamics and molecular associations in the nervous system. The line enables endogenous SNRNP70 to be visualised across nuclear, somatic and neuronal-process compartments. Moreover, this line can be readily combined with RNA labelling, cell-type-specific reporters, immunofluorescence and protein-proximity assays. Its localisation within dynamic neuronal mRNP granules provides an important application of the resource and demonstrates how endogenous tagging can reveal the spatial organisation of an RBP without reliance on overexpression. The TgKI(*snrnp70-eGFP*) line therefore provides a useful platform for future studies investigating how SNRNP70 and other RBPs contribute to spatial organisation and dynamics of RNA metabolism, while avoiding overexpression-associated artefacts.

## Acknowledgements

We thank the staff at the University of Bath Fish Facility and the University of Exeter Aquarium Research Centre for their excellent fish husbandry and care. We are also grateful to the staff of the Bioimaging Facilities at both universities for their technical support. We thank Stephanie Jones for her valuable comments on the manuscript. Finally, we thank Daniel Levic for kindly providing the eGFP knock-in construct and for his valuable advice.

## Author contributions

Conceptualisation: J.L-J., N.N. Data curation: J.L-J., N.N. Formal analysis: J.L-J., N.N. Funding acquisition: D.G., N.N. Investigation: J.L-J., S.S., N.N. Methodology: J.L-J., S.S., N.N. Project administration: N.N. Resources: J.L-J, D.G. Supervision: D.G., N.N. Writing – original draft: J.L-J, S.S. Writing – review and editing: J.L-J., S.S., D.G., N.N.

---

**Supplementary figure 1: Validation of eGFP insertion by gDNA sequencing**

Sequencing of wild-type (WT) and knock-in (KI) alleles in two separate animals. The start of the targeting cassette in intron 8 is indicated with a forward angle bracket. The gRNA target site is underlined. SNPs are highlighted in yellow and INDELs in red. The sequence highlighted in green corresponds to the targeting cassette and is present only in the KI allele. This sequence includes the base-pair substitution introduced to mutate the PAM site (immediately downstream of the gRNA target site), removal of the stop codon at the end of exon 10, and insertion of a small linker followed by eGFP immediately downstream of exon 10.

**Supplementary figure 2: RT-qPCR comparison of SNRNP70-eGFP KI and SNRNP70 WT transcript expression**

**(A)** RT-qPCR showing a comparison of gene expression between SNRNP70-eGFP KI and untagged transcripts in heterozygous F_1_ animals. Bar graph shows the gene expression ratios for the Exon10/eGFP (capturing the KI allele) and Exon10/3’UTR (capturing the wild-type allele), showing a non-significant difference in expression ratio between the KI and WT alleles. The graph shows mean values ± SEM. ns, not significant, Welch’s t-test.

**(B)** RT-qPCR showing the expression of the nearest neighbour protein coding gene (*lin7b*) in KI and *mitfa* control animals. The graph shows mean values ± SEM. ns, not significant, Welch’s t-test.

**Supplementary figure 3: ICC-IF of potential cytoplasmic SNRNP70 interactors**

**(A)** Representative image showing PABPC1B distribution within 12 DIV embryonic primary neurons. DAPI (blue); acetylated α-tubulin (red); and PABPC1B (magenta). Scale bar, 50 μm.

**(B)** Representative image showing FUS distribution within 12 DIV embryonic primary neurons. DAPI (blue); acetylated α-tubulin (red); and FUS (magenta). Scale bar, 50 μm.

**(C)** Representative image showing UPF1 distribution within 12 DIV embryonic primary neurons. DAPI (blue); acetylated α-tubulin (red); and UPF1 (magenta). Scale bar, 50 μm.

**Supplementary figure 4. SNRNP70-eGFP and PABPC1B localisation in the brain**

**(A1)** Anti-PABPC1B IHC-IF demonstrates broad expression of the protein throughout the forebrain, within both the telencephalon (T) and diencephalon (D) (encircled) of a 96 hpf larva. In addition, PABPC1B expression is visible within the inner plexiform layer (ipl) of the retina. Scale bar, 100 μm.

**(A2)** DAPI staining demonstrates that PABPC1B staining occurs both within the nuclei and cytoplasm of cells throughout the forebrain and retina. Scale bar, 100 μm.

**(A3)** Anti-GFP staining for SNRNP70-eGFP demonstrates widespread staining throughout the forebrain and retina. Scale bar, 100 μm.

**(A4)** Overlaying SNRNP70-eGFP staining with PABPC1B staining demonstrates broad co-localisation of both proteins within the telencephalon and diencephalon. In addition, both proteins show strong overlap within the inner plexiform layer (ipl). Furthermore, SNRNP70 shows strong expression within the optic nerve (arrow), from which PABPC1B appears to be largely absent. Scale bar, 100 μm.

**(A5)** Anti-acetylated α-tubulin staining confirms that SNRNP70 and PABPC1B are indeed co-localised within the neuronal processes of the retina, and confirms that SNRNP70 localises within the optic nerve (arrow). Scale bar, 100 μm.

**(A6)** Merged image of all channels: DAPI (blue); PABPC1B (red); SNRNP70-eGFP (green); acetylated α-tubulin (magenta). Scale bar, 100 μm.

**Supplementary figure 5. SNRNP70-eGFP and FUS localisation in the brain**

**(A1)** Anti-FUS IHC-IF demonstrates widespread, predominantly nuclear staining for FUS in the mid-hindbrain area of a 96 hpf larva. The region encircled corresponds to the myelencephalon, demonstrating widespread FUS localisation within this brain region. Scale bar, 100 μm.

**(A2)** Anti-FUS and anti-DAPI staining demonstrates that FUS is primarily localised within nuclei of the mid-hindbrain and myelencephalon. Scale bar, 100 μm

**(A3)** Anti-GFP staining for SNRNP70-eGFP demonstrates ubiquitous staining, as expected, along with a strong signal within the neuronal processes of the myelencephalon and mid-hindbrain. Scale bar, 100 μm.

**(A4-A5)** Anti-FUS/anti-GFP staining demonstrates broad co-localisation of SNRNP70 and FUS throughout the mid-hindbrain. Furthermore, continuous overlap of the two proteins is observed within the myelencephalon (upper left arrow, panels A4/A5), along with punctate overlap within neuronal processes (lower right arrow, panels A4/A5). Scale bar, 100 μm.

**(A6)** Merged image of all channels: DAPI (blue); FUS (red); SNRNP70-eGFP (green); and acetylated α-tubulin (magenta). Scale bar, 100 μm.

**Supplementary figure 6: SNRNP70-eGFP and UPF1 localisation in the brain**

**(A1)** Anti-UPF1 IHC-IF revealed widespread nuclear as well as cytoplasmic staining for the protein throughout the midbrain. Encircled are the regions inscribed by the tectum (Tec), tegmentum (Teg), and hypothalamus (Hyp), showing widespread presence of UPF1 in the first two brain regions, and weaker presence in the latter. Scale bar, 100 μm.

**(A2)** DAPI staining shows that UPF1 is predominantly nuclear throughout these brain regions, albeit with some cytoplasmic staining. Scale bar, 100 μm.

**(A3)** Anti-GFP staining shows ubiquitous SNRNP70-eGFP throughout the midbrain, with strong expression in the tectum and tegmentum. Scale bar, 100 μm.

**(A4)** Overlaying the UPF1 and SNRNP70 IHC-IF shows a broad degree of co-localisation, with both proteins present in the tectum and tegmentum. Scale bar, 100 μm.

**(A5)** Anti-acetylated α-tubulin IHC-IF shows that the SNRNP70-eGFP staining within the tectum is primarily cytoplasmic, as is the staining within the tegmentum. Scale bar, 100 μm.

**(A6)** Merge of all channels: DAPI (blue); UPF1 (red); SNRNP70-eGFP (green); acetylated α-tubulin (magenta). Scale bar, 100 μm.

**Supplementary video 1: SNRNP70^+^ RNP granule staying near the nucleus**

Representative example of an SNRNP70^+^ mRNP granule in the hindbrain at 48 hpf. The granule appears to be motile while remaining in close proximity to the nucleus throughout the duration of the video.

**Supplementary video 2: Motile SNRNP70^+^ RNP granule showing long-range movements**

Representative example of an SNRNP70^+^ mRNP granule in the hindbrain at 48 hpf. The granule exhibits long-range movements throughout the duration of the video.

