## Supplementary figures for "An SNRNP70-eGFP knock-in zebrafish line reveals the physiological localisation and dynamics of endogenous SNRNP70 during development"

### Supplementary figure 1: Validation of eGFP insertion by gDNA sequencing

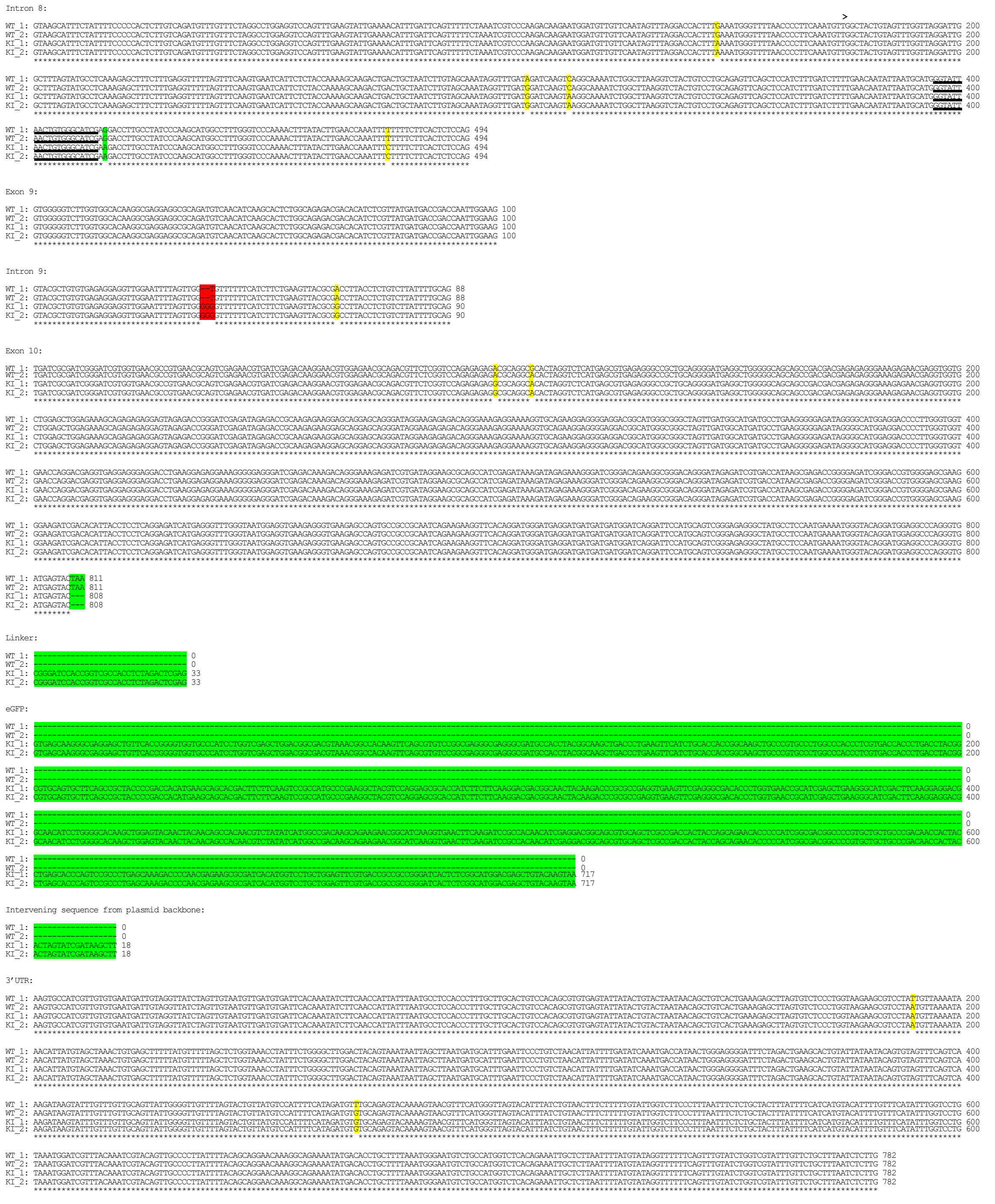


> start of the targeting cassette (320bp upstream to the 3' end of intron 8)
SNPs
INDELs
Sequence present within the targeting cassette
gRNA target site

**Supplementary figure 2: RT-qPCR comparison of SNRNP70-eGFP KI and SNRNP70 WT transcript expression**

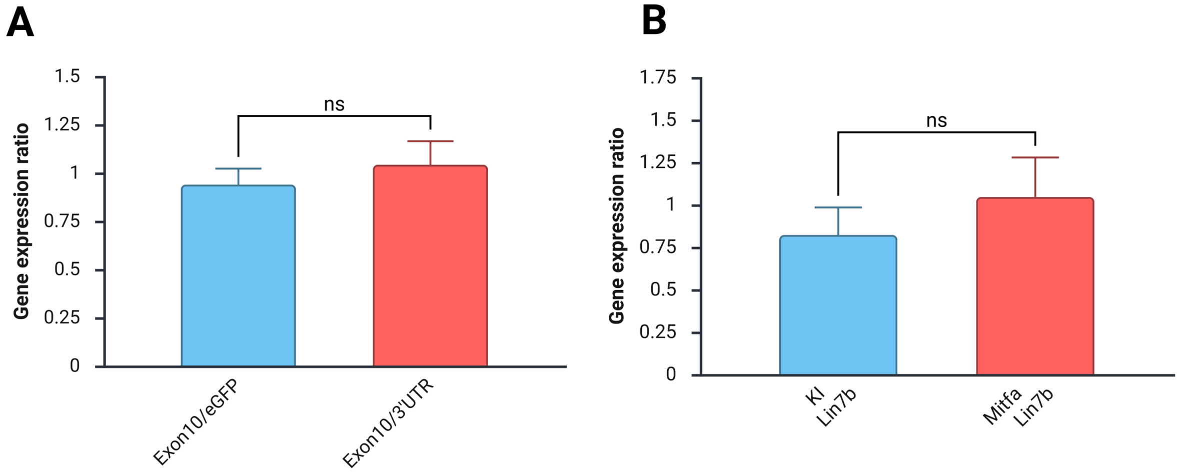

**Supplementary figure 3: ICC-IF of potential cytoplasmic SNRNP70 interactors.**

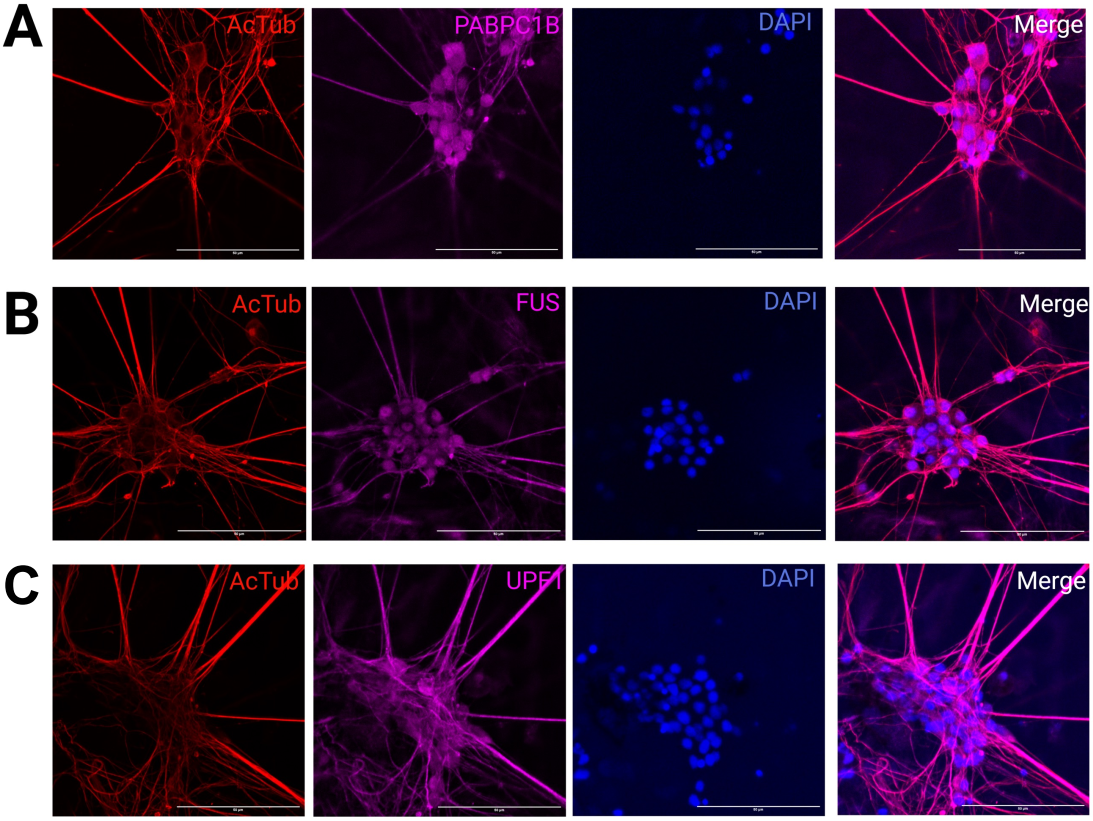

**Supplementary figure 4: SNRNP70-eGFP and PABPC1B localisation in the brain**

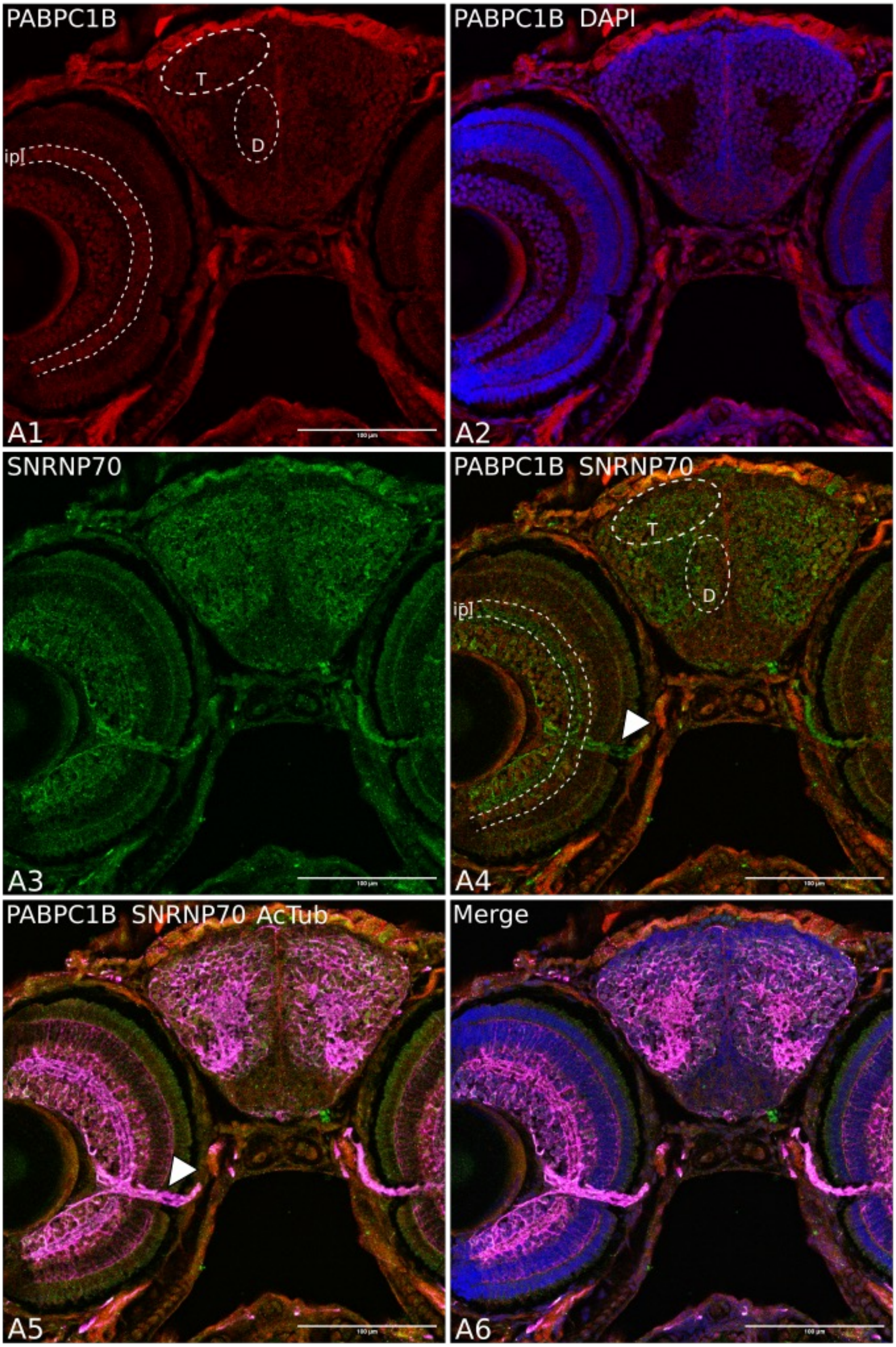

### Supplementary figure 5: SNRNP70-eGFP and FUS localisation in the brain

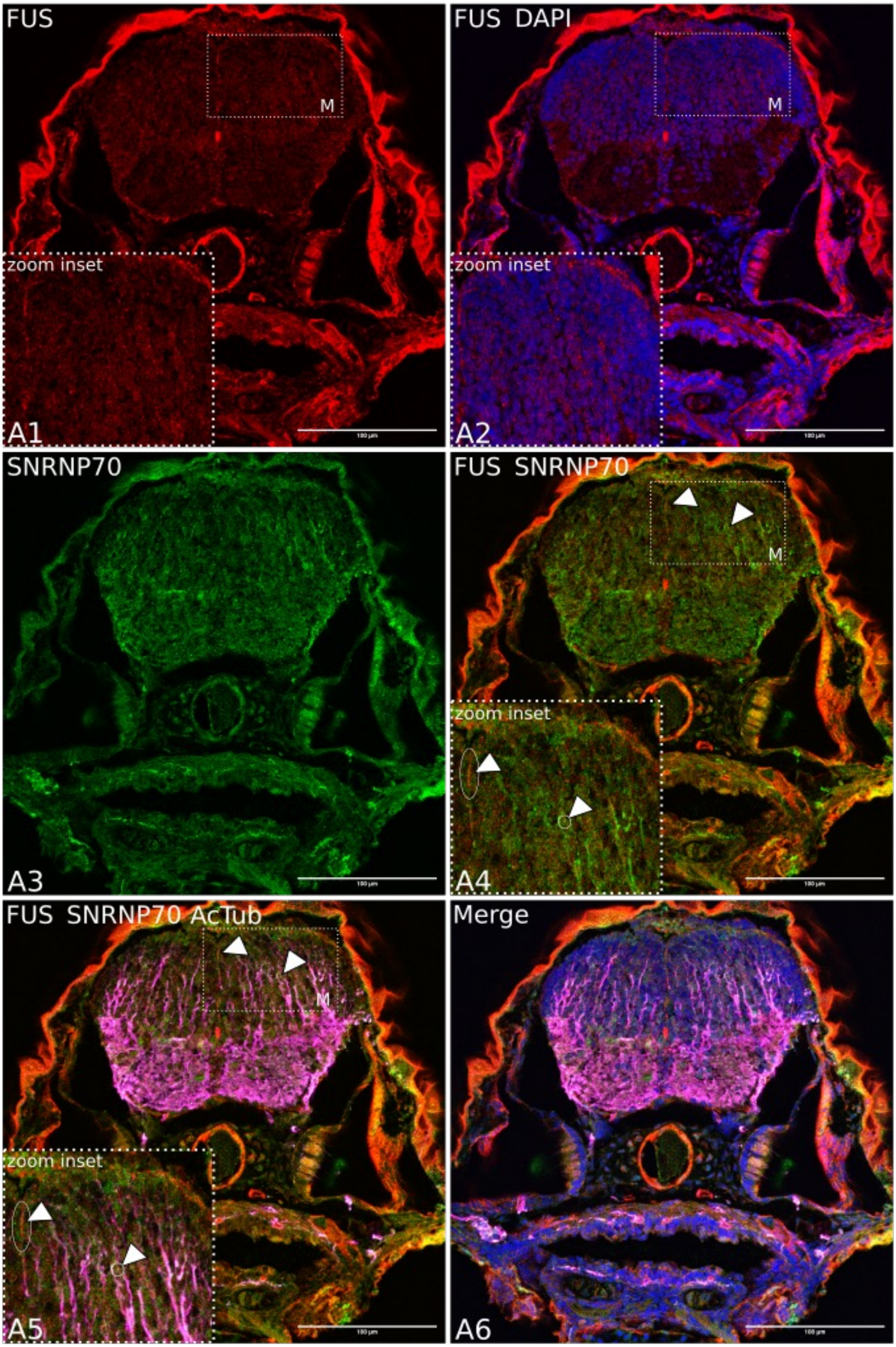

**Supplementary figure 6: SNRNP70-eGFP and UPF1 localisation in the brain**

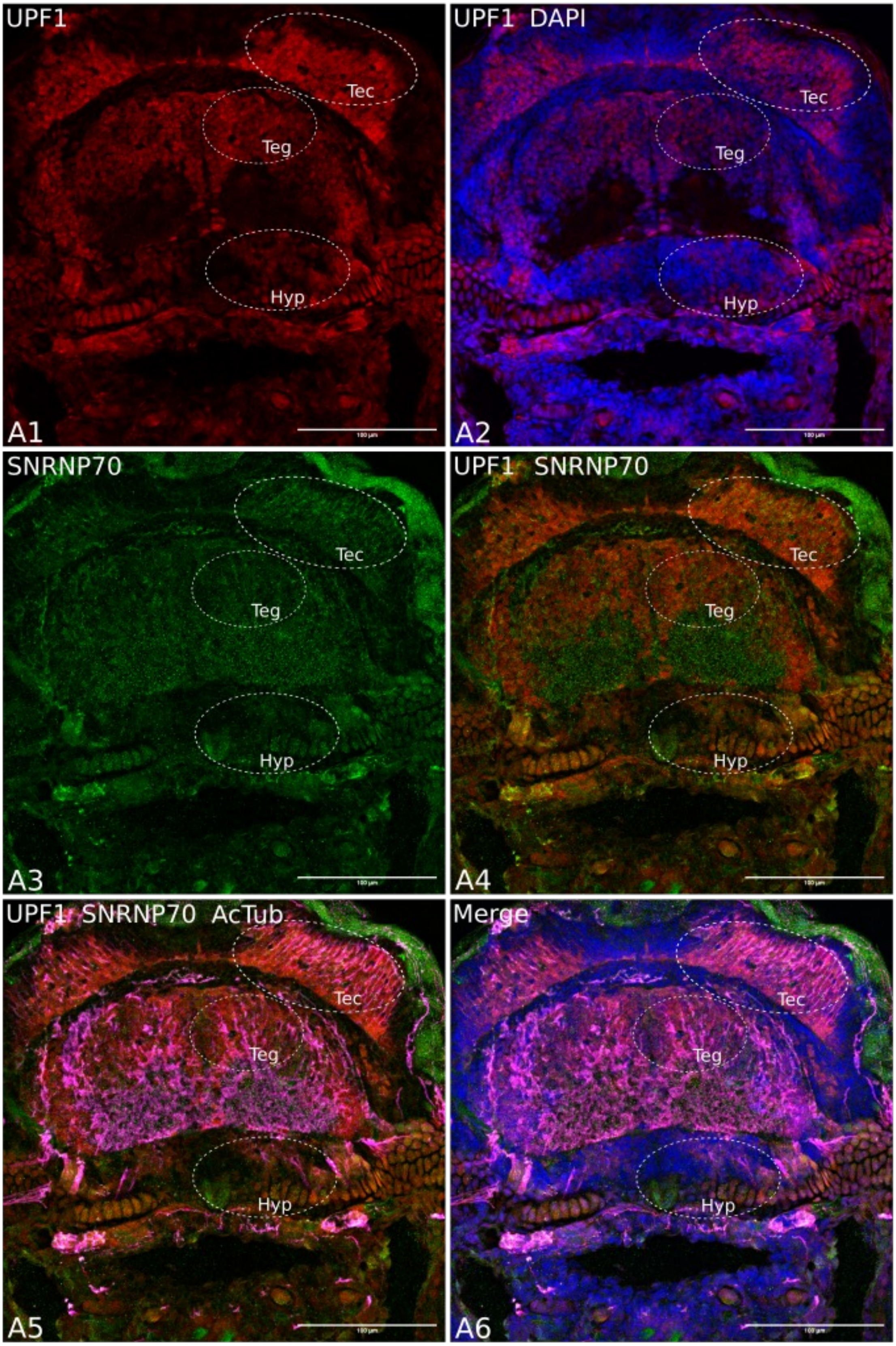
